# Sampling bias corrections for accurate neural measures of redundant, unique, and synergistic information

**DOI:** 10.1101/2024.06.04.597303

**Authors:** Loren Koçillari, Gabriel Matías Lorenz, Nicola Marie Engel, Marco Celotto, Sebastiano Curreli, Simone Blanco Malerba, Andreas K. Engel, Tommaso Fellin, Stefano Panzeri

**Affiliations:** Institute for Neural Information Processing, Center for Molecular Neurobiology, University Medical Center Hamburg-Eppendorf (UKE), Hamburg, Germany; Department of Neurophysiology and Pathophysiology, University Medical Center Hamburg-Eppendorf (UKE), Hamburg, Germany; Istituto Italiano di Tecnologia, Genova, Italy; Department of Pharmacy and Biotechnology, University of Bologna, Bologna, Italy

## Abstract

Shannon Information theory has long been a tool of choice to measure empirically how populations of neurons in the brain encode information about cognitive variables. Recently, Partial Information Decomposition (PID) has emerged as principled way to break down this information into components identifying not only the unique information carried by each neuron, but also whether relationships between neurons generate synergistic or redundant information. While it has been long recognized that Shannon information measures on neural activity suffer from a (mostly upward) limited sampling estimation bias, this issue has largely been ignored in the burgeoning field of PID analysis of neural activity. We used simulations to investigate the limited sampling bias of PID computed from discrete probabilities (suited to describe neural spiking activity). We found that PID suffers from a large bias that is uneven across components, with synergy by far the most biased. Using approximate analytical expansions, we found that the bias of synergy increases quadratically with the number of discrete responses of each neuron, whereas the bias of unique and redundant information increase only linearly or sub-linearly. Based on the understanding of the PID bias properties, we developed simple yet effective procedures that correct for the bias effectively, and that improve greatly the PID estimation with respect to current state-of-the-art procedures. We apply these PID bias correction procedures to datasets of 53117 pairs neurons in auditory cortex, posterior parietal cortex and hippocampus of mice performing cognitive tasks, deriving precise estimates and bounds of how synergy and redundancy vary across these brain regions.

## 1 Introduction

It is widely believed that the brain is a complex system and that behavior emerges from the organized pattern of the interactions between the brain’s computing elements - the neurons [1].

Because of this, the study of how interactions between neurons shape information processing has fascinated computational and empirical neuroscientists for decades [2, 3, 4, 5, 6]. Information theory has been a prominent tool in this research as it is uniquely positioned and is a natural choice for investigating neural information processing [7, 8, 9]. Shannon information captures all ways in which systems carry information, it is highly general and applicable regardless of the type of noise and statistics, and is thus equally applicable across species and recording modalities and to real data and in silico models. While earlier work has concentrated on understanding whether correlations between the activity of different neurons increase or decrease information [10, 3, 4, 11], recent advances in information theory based on Partial Information Decomposition (PID) [12] have enabled neuroscientists to formulate more precise questions [13] about the unique information carried by each individual neuron, the synergistic information generate by neural interactions (that is, new information that emerges specifically from and is found only in the interactions between neurons) or redundant information present when different neurons share the same information.

Earlier applications of Information Theory to neuroscience have recognized that Shannon Information measures from neural activity suffer from a prominent limited sampling bias [14, 15, 16]. This bias does not only affect the precision of the estimate, but can also skew comparisons between the information carried by simpler (less biased) and more complex (more biased) neural representations. The information theoretic neuroscience literature has provided effective tools for correcting for the Shannon information sampling bias [17]. However, to date the problem of whether synergy, redundancy and unique information components of PID are biased has not been studied systematically. To our knowledge, only one study investigated PID sampling bias [18]. This method was valid only for Gaussian distributions, which do not apply easily to the discrete spiking activity of neurons. Here, we study the sampling bias of discrete PID estimators suitable to study the spiking activity of neurons. We found that the PID components are unevenly biased, with synergy far more biased than any other component. We study and provide an understanding of the origin and properties of the bias, and based on this we provide methods to reduce this problem effectively that provide more accurate measures than the state of the art. Finally, we test and apply our method to 53117 pairs of neurons simultaneously recorded from the mouse cortex and the hippocampus.

## 2 Background: short introduction to PID

For completeness, we first summarize the concepts of PID that we use. PID decomposes the information jointly carried by a set of source variables (for us, a set of simultaneously recorded neurons) about a target S (for us, a sensory stimulus) into non-negative components that capture information about the target that is either redundantly encoded across sources, uniquely encoded by a single source or synergistically encoded by the combination of sources. In this paper, we will focus on the case of two source variables, which has received the most attention in the PID literature and in real data applications, and which has the most established theoretical foundations [19, 20, 21, 18, 22, 23, 24]. We focus our presentation on the information about sensory stimuli carried by neurons, but this framework straightforwardly extends to information carried by neurons about other quantities, such as cognitive or motor variables or the activity of other neurons. We consider the Shannon information *I*(*S*; *R*_1_, *R*_2_) about an external stimulus *S* carried jointly by the neural spiking activity *R*_1_ and *R*_2_ of two simultaneously recorded neurons and the Shannon information *I*(*S*; *R*_*i*_) that each of the two neurons (*i* = 1, 2) carries about *S. I*(*S*; *R*_1_, *R*_2_) and *I*(*S*; *R*_*i*_) are computed from the probability distributions *p*(*S, R*_1_, *R*_2_) and *p*(*S, R*_*i*_) (*i* = 1, 2) [25, 26], as follows:

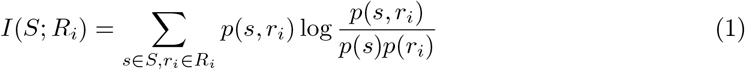

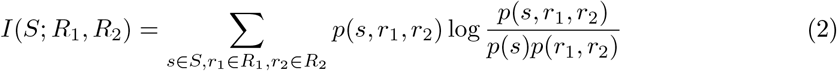

PID decomposes the joint and single-neuron information into four non-negative components that satisfy the following linear relationships [12]:

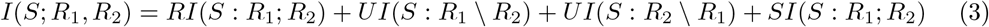

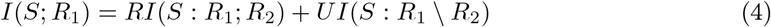

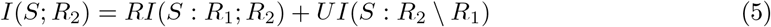

where in the above equations *RI*(*S* : *R*_1_; *R*_2_) is the redundant (or shared) information that both *R*_1_ and *R*_2_ encode about *S*; *UI*(*S* : *R*_1_ \ *R*_2_) and *UI*(*S* : *R*_2_ \ *R*_1_) are the unique information about *S* provided by one neuron but not by the other; and *SI*(*S* : *R*_1_; *R*_2_) is the synergistic information about *S* encoded by the combination of *R*_1_ and *R*_2_. (We will sometimes shorthand these components as 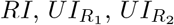, and *SI*.) Eq. (3-5) mean that all 4 components contribute to the joint information, whereas only the information that a source carries uniquely and the information that is redundantly carried by both contribute to single-neuron information. Because the 4 PID components satisfy 3 linear constraints (Eqs. 3-5), determining one component is sufficient to compute the other three. We provide explicit equations of all components as function of synergy in Eqs (S12).

Several definitions of PID components have been proposed [27, 28], satisfying desired properties including non-negativity of each component and symmetry of *RI* and *SI* under permutation of *R*_1_, *R*_2_. We will mostly use the BROJA definition [20], as it satisfies many desirable properties including additivity of *RI, SI*, and *UI* for independent systems of sources and targets [28, 29] and has been extensively applied to neural data [23, 30, 24, 31]. The BROJA defines the Union information, that is the target information in the joint source space that cannot be possibly attributed to synergistic interactions, or equivalently the total target information that can be extracted from a single source, as:

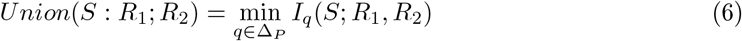

where Δ_*P*_ is the set of all joint probability distributions *q*(*S, R*_1_, *R*_2_) that have the same pairwise marginals *q*(*S, R*_1_) = *p*(*S, R*_1_) and *q*(*S, R*_2_) = *p*(*S, R*_2_) as the original distribution *p*(*S, R*_1_, *R*_2_), and *I*_*q*_(*S*; *R*_1_, *R*_2_) is the joint information computed for distribution *q*(*S*; *R*_1_, *R*_2_). Then, the synergy is defined as the difference between the joint and the union information:

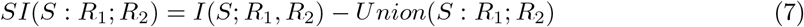

Other PID definitions, which were also successfully applied to neuroscience [22, 32, 33] but do not satisfy additivity include the originally proposed *I*_*min*_ [12] and the minimum mutual information *I*_*MMI*_ [21]. *I*_*min*_ quantifies *RI* as the similarity between *R*_1_ and *R*_2_ in discriminating individual values of *S*, while *I*_*MMI*_ quantifies *RI* as the minimum between the mutual information individually carried by *R*_1_ and *R*_2_, thus capturing only the amount but not the content of information carried by each neuron [28].

## 3 Background: discrete estimators of information and PID in neuroscience

Neural spiking activity is an intrinsically discrete variable. Thus, in most information theoretic studies of neural spiking activity, the responses have been treated as discrete variables. When focusing on spike count codes, the neural response is described by the number of spikes emitted by the neuron in the time window of interest [34, 35, 36, 37, 38, 39, 40]. When investigating if the timing of spikes encodes additional information above and beyond that present in the spike counts, then the most established approach [41, 42, 43, 44, 45] is to discretize the neural response time window of interest into a number of small time bins and then turn the spike train into a binary word. Importantly, often neural responses are sparse and information is encoded by relatively low spike numbers [36, 46]. In such cases, as we show in Fig. S12, the Gaussian approximation to the information is highly inaccurate. Because it is simple and does not require assumptions on the probability distributions, discretization of neural responses for computing information has been used to compute information also from continuous-valued non-spiking aggregate measures of neural activity such as LFP, EEG, or fMRI [47, 48, 49, 50]. In addition, often only a discrete number of different stimulus conditions is presented in an experiment, and thus the stimulus *S* is typically a discrete variable. Under such conditions, the information theoretic quantities can be computed by simply estimating the probabilities by the empirical occurrences across experimental trials (maximum likelihood estimators) and plugging them into the information equations. This discrete information approach, which we call the plugin approach, has been extensively used in neuroscience for both Shannon information [41, 42, 43, 35, 3, 51, 37, 38, 52, 53, 54, 39, 55, 40, 31] and PID [56, 57, 58, 53, 32, 59, 23, 22, 33, 30, 31, 50]. This discrete PID approach has also been used extensively across fields of biology and applied sciences [60, 61, 62, 63].

## 4 Numerical investigation of the bias of individual PID components for discrete estimators

Calculation of information requires accurate estimation of the stimulus-response probabilities. With an infinite amount of data, the true stimulus-response probabilities could be measured exactly. However, any real experiment only yields a finite number of trials from which probabilities must be estimated. The estimated probabilities have finite sampling fluctuations around their true values (Fig. S2) which lead to both systematic error (bias) and statistical error (variance) in estimates of information (Fig. S2 and Supplemental Material, SM Section SM1.3). While variance can be reduced by averaging (e.g. across experimental subjects or groups of neurons), the bias cannot.

The sampling bias properties of discrete estimators have been extensively studied for Shannon information [14, 15, 17], but not to our knowledge for discrete PID estimators. To document them for PID, we simulated spike count responses of a pair of individual neurons in response to a set of *S* = 4 different stimuli. We studied how the estimate of each PID component depends on the number of available trials. We focus the presentation on the BROJA PID decomposition [20], but we confirm in Fig. S6, S7, S10, S11 that similar results apply to *I*_*min*_ and minimum mutual information *I*_*MMI*_.

We developed three different scenarios with varying degrees of synergy and redundancy. In each case, the spike count *r*_*i*_ (*i* = 1, 2) of each of the two simulated neurons for each stimulus was the sum of two Poisson processes. One Poisson process, that was independently drawn for each neuron, expressed the variability of responses “private” to each neuron. Another Poisson process was shared between the two neurons and gave rise to across-neuron correlations. Within each scenario, we also varied the overall level of information, because previous studies showed that the bias of Shannon information depends on it [17]. In our simulations, the parameter *B* expressed the baseline level of activity; parameter *α* regulated the strength of stimulus tuning of each neuron (increasing *α* increased single-neuron information); parameter *β* regulated the dissimilarity of tuning between neurons (smaller *β* meaning more independent tuning); a parameter *γ* regulated the strength of the shared process. The overall information level was increased by increasing *α* or *γ* or reducing *B*. Redundancy was increased by increasing *β*. Synergy was increased by increasing *γ*.

In Fig. 1 we plot the values of the plugin estimates of joint information and of the PID terms as function of the simulated number of trials per stimulus, averaged over all *n* = 96 repetitions of the simulations with the considered number of trials. In these simulations we discretized the spike counts of each neuron into 4 equipopulated bins (leading to 16 possible discrete joint responses).

**Figure 1.**
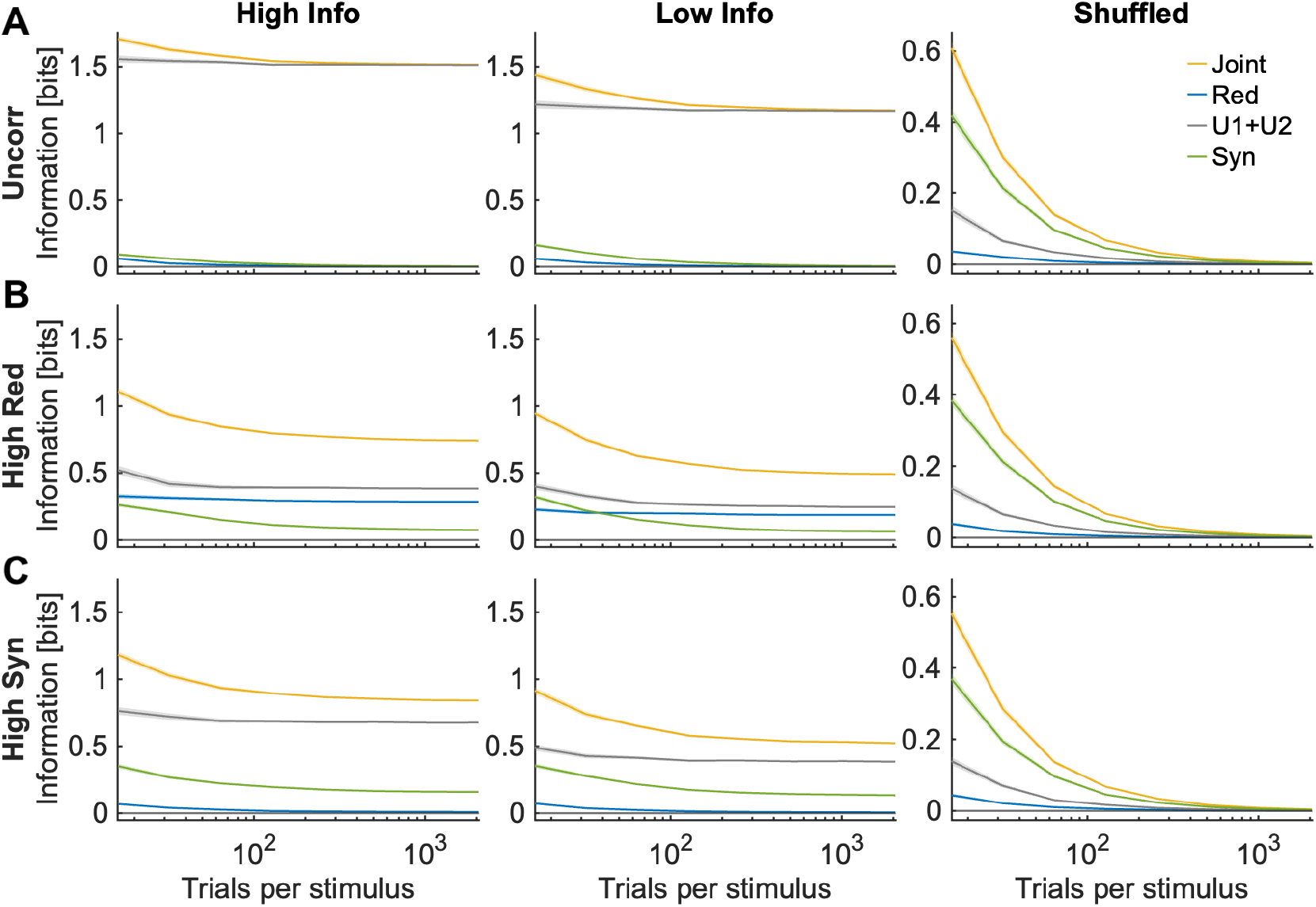
Joint information and PID quantities as a function of the number of simulated trials used to compute them. Top, central and bottom rows plot the simulated scenarios with no interaction, high redundancy and high synergy, respectively (see SM Section SM1.5). Left, center and right columns represent simulations with higher information (*α* = 10), lower information (*α* = 7) and with shuffled low-information data. “Syn”: synergy. “Red”: redundancy. “U1+U2”: sum of the two unique information of each neuron. Here we used the plugin method without bias corrections. We used *R* = 4 discretization bins for each neuron (Table S1). Each panel plots mean ± 2 SEM over *n* = 96 simulations.

The limited sampling bias in the information estimates can be visualized by comparing the average value of information obtained with a given number of trials with the asymptotic value obtained with the largest number of trials. In all scenarios, the plugin estimate of the joint Shannon information was biased upward and the bias decreased with the number of trials, as reported previously [17, 43]. Here, we focus on the bias of the estimates of the PID quantities. We found several highly consistent and important results. First, as for Shannon joint information, PID quantities were biased upward, with the bias decreasing smoothly with the number of trials. Second, for the tens of trials usually available in real experiments, the bias could be as large, or even larger, than the target information quantities, meaning that the bias must be corrected for real data analyses. Third, and unlike what was assumed in previous studies [18], the bias is highly uneven across PID components. The synergy was by far the PID component with the largest upward bias. Its bias was lower than but comparable to that of the joint information. Unique information was also biased upward, albeit much less so than the synergy. Redundancy was almost unbiased. This is important because it shows that conclusions taken from limited empirical data without considering the bias will produce estimates artificially biased toward synergy. For example, in simulations with ground-truth values of redundancy larger than synergy, we would have incorrectly estimated synergy larger than redundancy for lower number of trials due to the sampling bias. Fourth, the bias was larger, both proportionally and in absolute terms, for lower information levels.

In SM Section SM1.8 we report analytical approximations to the sampling bias that allow an intuitive understanding and support the generality of these findings. The main take-home message from these calculations is as follows. For large enough numbers of trials, the bias can be expanded in inverse powers of the number of trials *N*, accounting for the smoothly decreasing bias with the number of simulated trials. The spurious levels (upward bias) of information is due to the fact that random fluctuations in probabilities make the probabilities more different across stimuli than they actually are. At fixed numbers of trials, the bias induced by these fluctuations depends only on the number of possible responses and the bias is larger for larger number of discretized possible responses. The main term for the synergy bias is due, like for the joint information, to fluctuations in the joint probability *P* (*r*_1_, *r*_2_, *s*), which is more undersampled than the marginal probabilities. The main bias term for the unique information originates from and is explained by fluctuations in the marginal probabilities, which are smaller because the marginal probabilities are defined in a smaller single-neuron space and are easier to sample. As a result, the bias of the joint information and of the synergy increases quadratically with the number of single-neuron discrete responses, whereas the bias of the unique and redundant information increase linearly or sublinearly (as found numerically, compare Fig. 1, S4, S5). Low information levels typically correspond to stimulus-specific distributions with a larger number of possible responses.

## 5 Correcting for the PID limited-sampling bias

Having documented and understood the properties of the bias of the PID, we now use this knowledge to propose and evaluate methods for bias correction.

Because the PID bias decreases smoothly with the number of trials, and because analytical calculations show that when the number of trials is sufficiently large, the bias depends polynomially on the inverse of the number of trials 1*/N*, we extend the Quadratic Extrapolation (QE) procedure originally proposed in Ref [43] for Shannon information to correct for the bias of each PID term. We recomputed plugin PID terms using half and a quarter of the available data, we fitted these information values to a polynomial in 1*/N* and we used the best-fit coefficient to estimate and remove the bias. The QE bias correction substantially improved the estimates of all PID quantities. While plugin bias-uncorrected estimates of synergy needed large numbers of trials per stimulus (*N*_*s*_ ≈ 512 − 1024 trials for 16 joint response bins (Fig. 1), corresponding to 32-64 trials per stimulus and joint response bin when *R* is varied, (Fig. S4, S5)), the QE reached accurate estimates with small residual bias with almost an order of magnitude less trials (*N*_*s*_ ≈ 64 − 128 trials for 16 joint response bins, corresponding to 4-8 trials per stimulus and joint response bin (Fig. 2, 3, S8, S9)). The QE-corrected PID values are relatively accurate but not conservative because they have an upward residual bias (because information bias terms of higher order in 1*/N*, not fitted in the QE, are all positive [64]).

**Figure 2.**
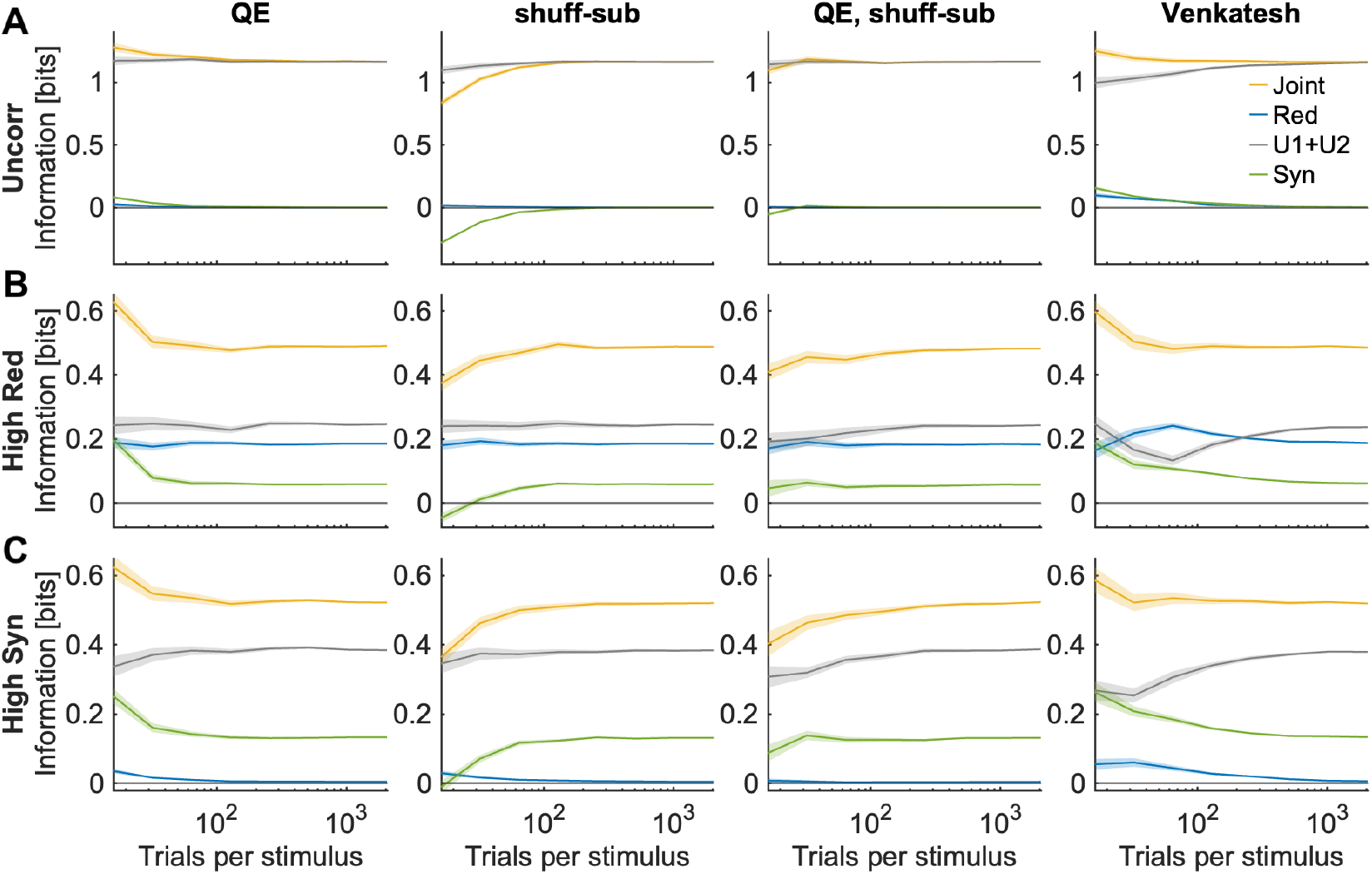
Performance of bias corrections with 4 discretization bins for each neuron. Joint information and PID quantities as a function of the number of simulated trials used to compute them. Top, central and bottom rows plot the simulated scenario with no interaction, high redundancy and high synergy, respectively (see SM Section SM1.5). Left to right columns report results of the QE, shuffle-subtraction, QE with shuffle-subtraction, and Venkatesh procedures, respectively. In each panel we plot the mean ± 2 SEM over *n* = 96 simulations.

**Figure 3.**
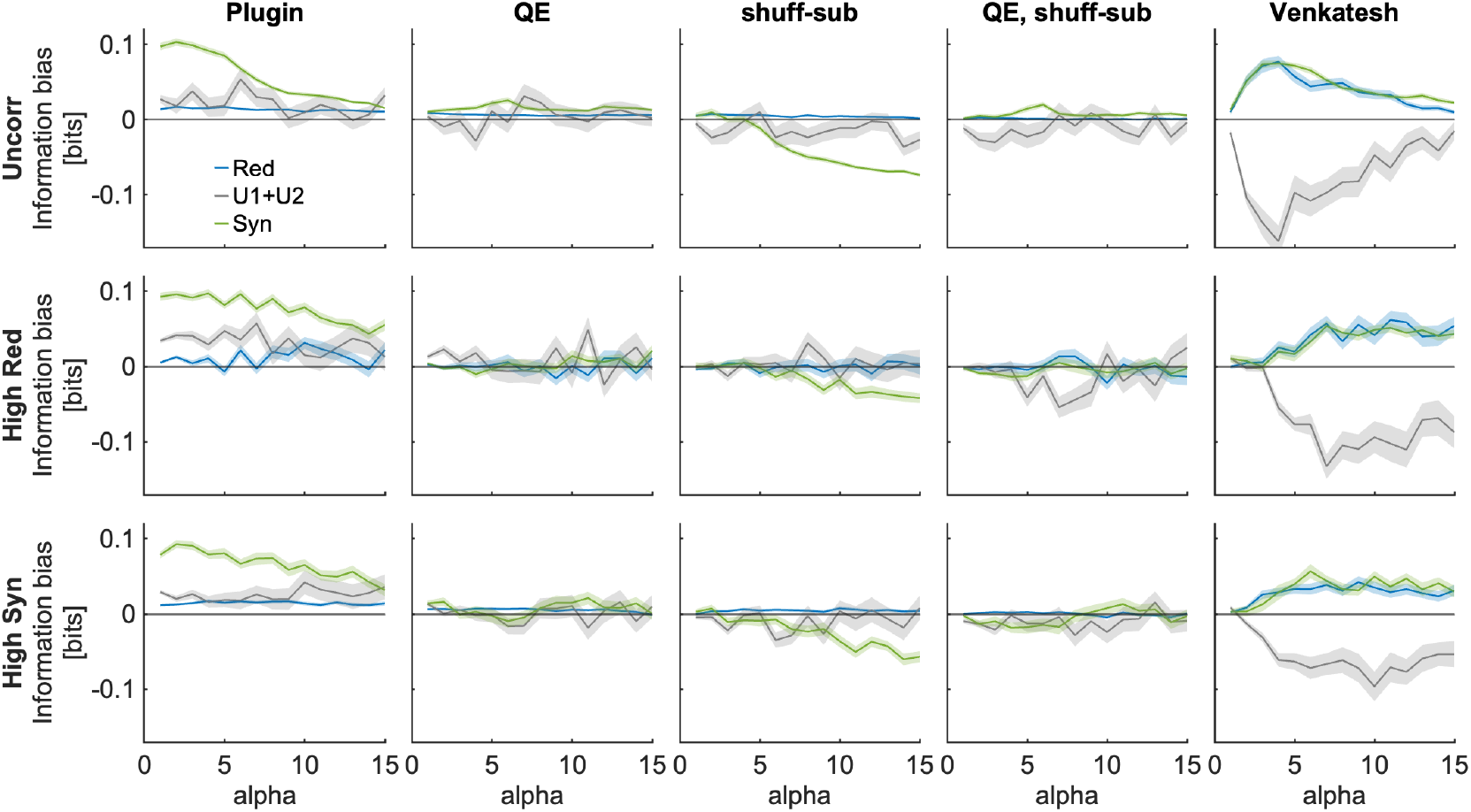
Performance of bias corrections using *N*_*s*_ = 64 trials per stimulus with 4 discretization bins for each neuron. In each panel we plot (rather than the information component value itself) the information component bias (computed as the information component estimated with the considered number of simulated trials minus the asymptotic information component estimated using the largest available number of simulated trials, that is 2048 trials per stimulus) as a function of the parameter *α* increasing single-neuron information in the simulated data. Top, central and bottom rows plot the simulated scenario with no interaction, high redundancy and high synergy, respectively (see SM Section SM1.5). Left to right columns report results with plugin estimators and with the QE, shuffle-subtraction, QE with shuffle-subtraction, and Venkatesh procedures, respectively. In each panel we plot mean ± 2 SEM over *n* = 96 simulations.

We then introduced a second PID bias correction (“shuffle-subtraction” bias correction) that subtracts from the plugin value of each term the value of the same term obtained after randomly shuffling the stimulus-response association. In the shuffled data, all information about the target (the stimulus) is destroyed. Thus, the plugin shuffled PID values can be taken as a bias estimate because they should be zero for infinite trials. Using this correction also increased the precision of the estimates compared to the plugin values (Fig. 2, S8). Importantly, and as supported by the analytical bias expansion (SM1.8), because lower information levels have larger upward bias, we observed a higher bias in the shuffled data than in the corresponding unshuffled data (Fig. 1). Thus, the shuffle-subtraction correction provides conservative estimates and can be used to lower-bound the PID estimates.

Given that the QE and shuffle-subtraction provide positive and negative residual PID bias respectively, we introduced a third bias correction procedure (“QE with shuffle-subtraction”) which combines the two operations. This procedure gave highly accurate results over the entire range of trials tested. Because the shuffle-subtraction provides conservative estimates, this third procedure was more conservative than the pure QE and it gave most often (though not always) negative residual bias.

Importantly, when there were enough trials for the bias corrections to work well (*N*_*s*_ ≈ 64 −128 trials for a joint distribution for 16 response bins, corresponding to 4-8 trials per stimulus and joint response bin when *R* is varied, see Fig. 2, 3, S8, S9), the QE and the QE with shuffle-subtraction gave nearly identical estimates and the pure shuffle-subtraction gave only slightly conservative and accurate estimates, suggesting that comparing on real data several bias correction methods on the same data may be beneficial for gaining confidence on the accuracy of the estimates.

The only bias correction which was proposed so far for PID was the one we term Venkatesh correction proposed in Ref [18]. It assumes that the Union information has the same bias as the joint information. Then it rescales all PID accordingly and implements post-hoc rectifications to make sure the PID terms are non-negative and still respect the PID properties of Eq. (3-5). Because, as we demonstrated above, the union information is much less biased upward than the joint information, this procedure leads to a major underestimation of union information, which then leads to major overestimations of synergy and redundancy that are present also for relatively large numbers of trials. We thus do not consider this correction further.

There are other bias corrections in the neural literature of discrete estimation of Shannon information [14, 65, 16, 15]. We did not consider them here because their derivation has not been extended to PID and because they performed worse than or equal to the QE with discrete estimators of Shannon information when tested on realistic simulations of neural population activity [17].

Although in the above we simulated *S* as a sensory stimulus, in many neuroscience applications *S* is the activity of other neurons. Our asymptotic expansions and considerations are valid as long as *S* is a discrete variable. In SM Section SM1.11 and Fig. S13 we show that the bias exists also when the target variable *S* is the discretized activity of another neuron and we show that the bias corrections are effective also in that case.

## 6 Evaluation of bias correction procedures on real neural data

We evaluated the utility of the PID bias correction using 3 datasets from previously published studies recording simultaneously with two-photon calcium imaging the activity of many neurons from the brain of mice performing cognitive tasks.

The first dataset consisted of *n* = 6209 pairs of neurons simultaneously recorded from auditory cortex during a sound intensity discrimination task [23]. We computed the information that the neurons carry about the sound intensity (a binary *S* = 2 stimulus set consisting of high vs low tone intensity). The activity of each neuron was first deconvolved to estimate the time-localized spiking activity from the calcium fluorescence signal imaged from each neuron. To compute the neural response variables *r*_1_ and *r*_2_ that enter the information calculations, for each neuron we summed the deconvolved activity within a 333-ms time window centered around the time of maximal information, and we discretized this signal into *R* = 3 bins (see SM Section SM1.10 for full details).

The second dataset [53] consisted of *n* = 10750 pairs of neurons simultaneously recorded from posterior parietal cortex (PPC) during a sound localization task in which mice reported perceptual decisions about the location (left or right of the midline) of an auditory stimulus while navigating through a visual virtual reality T-maze (Fig. 4B). Because PPC is an area involved in converting sensory information into perceptual decisions, we computed the information that the neurons carry about whether the sound came from left or right of the midline (*S* = 2 stimuli), corresponding to the sound location categorization that the mouse had to perform to turn toward the reward location. To compute the neural response variables *r*_1_ and *r*_2_ used for information calculations, for each neuron we summed the deconvolved activity within 320 ms time windows centered around the time of maximal information and we discretized this signal into *R* = 3 bins (see SM Section SM1.10).

**Figure 4.**
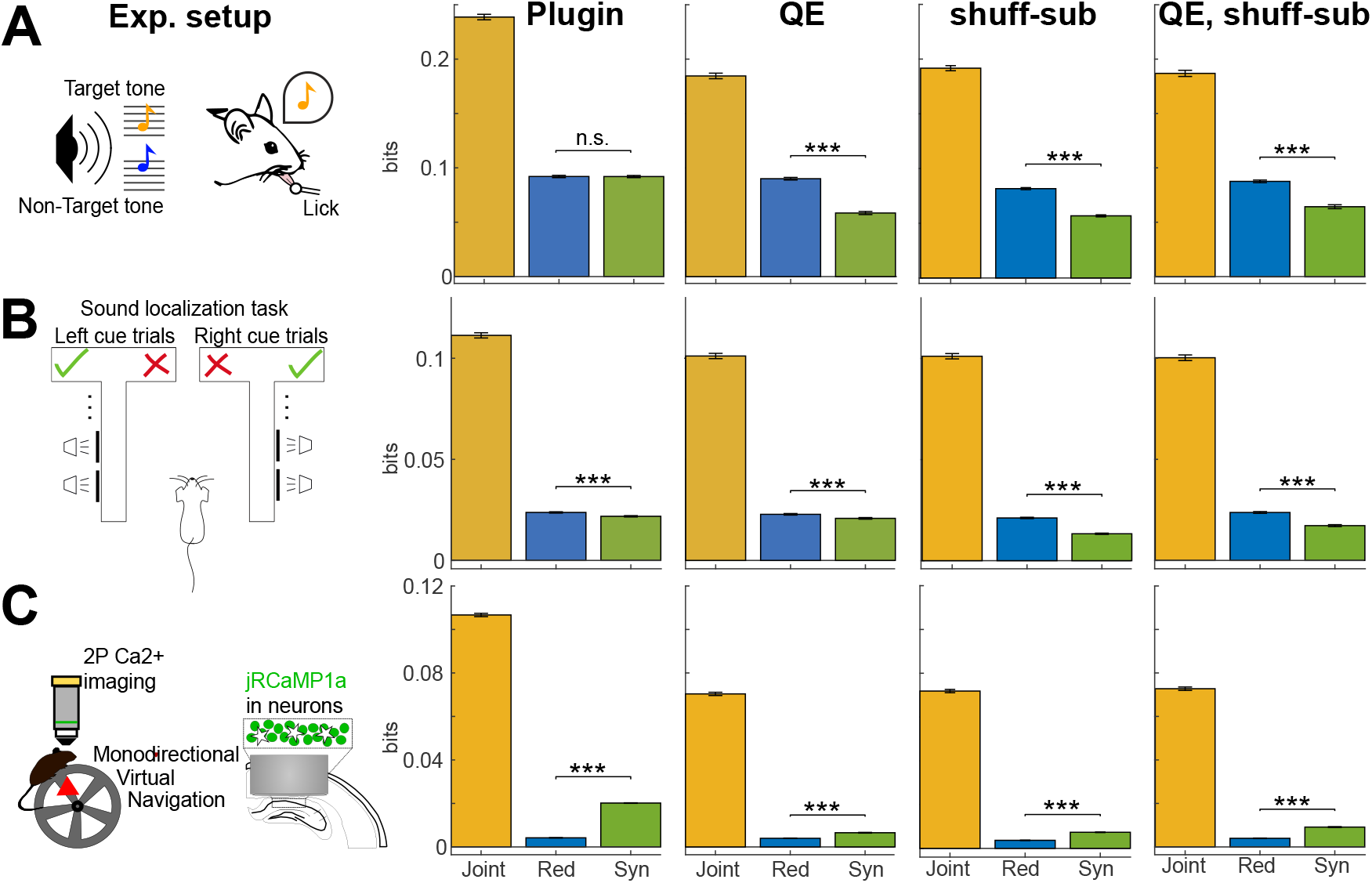
PID bias corrections on real neural data. Each panel plots mean ± 2 SEM over all analyzed simultaneously recorded neural pairs (*n* = 6209, 10750, 36158 for auditory cortex (top row), posterior parietal cortex (middle) and hippocampus (bottom) of joint information, synergy and Redundancy. Mean number *N*_*s*_ of available trials per stimulus per dataset was 70, 100, and 72, respectively. Columns from left to right plot: schematic of each task; results with plugin, QE, shuffle-subtraction, QE with shuffle subtraction respectively. Comparisons between synergy and redundancy were performed with a two-tailed paired t-test (^*∗∗∗*^*p <* 0.001, n.s.:*p >* 0.05).

The third dataset consisted of *n* = 36158 pairs of neurons simultaneously recorded from the CA1 region of the hippocampus [55] while mice navigated a linear track in virtual reality (Fig. 4B). Because the hippocampus encodes position in space, we computed the information that the neurons carry about the spatial location along the linear track (the location was discretized in *S* = 12 spatial bins). The neural response variables *r*_1_ and *r*_2_ used to compute information were the activity of the neurons in one imaging frame (333 ms) when the mouse was in a given position. As the slow kinetics of the calcium indicator and the slower imaging frame rate made it difficult to deconvolve the calcium fluorescence traces to estimate spiking activity, following Ref. [55] we discretized the *δF/F* calcium traces into *R* = 2 equipopulated bins (low and high activity) (see SM Section SM1.10).

We computed the joint information between the above-defined activity *r*_1_, *r*_2_ of simultaneously recorded pairs of neurons and the above defined stimulus *s*. We broke up this information into PID components using BROJA PID [20, 66]. We computed all 4 PID terms in Eq. (3). However, we focus the analysis and the neuroscientific interpretation on the differences between synergy and redundancy, to understand how these two emergent properties shape neural population coding in different brain regions. To test and exemplify the use of the bias-corrected PID algorithms, we compared (Fig. 4) the plugin estimates of PID with those obtained after applying the 3 bias corrections that we developed.

As shown by the reduced values obtained after applying the bias corrections, the plugin joint information and synergy were biased upward (consistent with simulations and theory). Despite the good number of trials available in these experiments, the synergy bias was substantial. Using the uncorrected plugin estimator would have led to a considerable overestimation of the bias and to a qualitative change of results in two datasets. In the auditory dataset (Fig. 4A), because of the large synergy bias, the uncorrected plugin estimator could not detect a significant difference between synergy and redundancy, whereas all 3 bias-corrected estimates consistently detected with high significance that redundancy was higher than synergy. In the hippocampal dataset (Fig. 4B), both the plugin and the 3 bias-corrected estimates reported a higher level of synergy than redundancy. However, the use of the plugin bias-uncorrected method would have overestimated by 211% the amount of synergy with respect to what was consistently found with the bias corrections. The PPC data (Fig. 4B) gave consistent values of higher redundancy than synergy with all methods.

Another important result is that, reassuringly, we found highly consistent estimates of both redundancy and synergy across bias correction methods. Redundancy was essentially identical across methods for each dataset. Synergy estimates obtained with QE and QE with shuffle subtraction were within 3% of each other in each dataset, suggesting that these estimates on real data are precise and unbiased.

The more conservative shuffle subtraction underestimated (only slightly for the auditory and hippocampal dataset, and a little more for the PPC dataset) the less conservative synergy values obtained with QE or QE with shuffle subtraction. However, and importantly, the fact that the synergy obtained with the conservative shuffle subtraction is positive proves that there is genuine synergy that cannot be due to finite sampling artifacts.

## 7 Discussion

We found that PID components suffer from a considerably limited sampling bias. Under simulated conditions relevant to neuroscience experiments, the bias was as large as the information quantities to be estimated, and thus cannot be neglected in neuroscience applications. Importantly, our work highlights and explains the presence of a major and previously neglected difference across PID terms in the sampling bias, which inflates synergy disproportionately. Neural synergy has been widely reported in recent years and has led neuroscientists to rethink how the brain integrates information. Our discovery calls for a careful re-evaluation of these reports with bias-corrected estimator.

We provided an analytical understanding of the properties and origin of the bias in terms of simple properties of experimental design (number of trials) and of analysis setting (number of discrete or discretized responses). Thus, our work helps informing experimental design.

Importantly, we provide generally applicable algorithms that correct for the bias and greatly improve information estimates with respect to state-of-the-art neural measures, which either neglected the bias problem (using uncorrected estimators) or correct for the bias using the incorrect assumptions that the bias is even across components. Importantly, the algorithms we develop not only improve in a major way the estimates, but some of these algorithms present positive and some other present negative residual estimation errors, allowing to empirically bound estimates with reasonable confidence.

From the neuroscientific point of view, applications of our method to simultaneous recordings with cellular resolution, confirmed the usefulness of the method to obtain reliable conclusions, highlight a widespread presence at the cellular level of synergy and redundancy in neuron-to-neuron interactions, and of region-to-region variations of the relationship between synergy and redundancy which were previously reported only at the level of aggregate signals without cellular resolution [32, 13].

### Limitations

Limitations of this work that require further theory and simulations include (1) We tested the bias of BROJA, *I*_*min*_ and *I*_*MMI*_ PID definitions. It would be important to test others. (2) The bias correction procedures are heuristic, although we confirm and support them by providing an analytical expansion of the bias derived in the large *N* limit. However, our derivation is partly heuristic and we did not provide theoretical guarantees of sign of residual errors of different bias correction procedures. (3) We tested information between pairs of neurons and stimuli but we have not tested large populations. The direct calculation of information is very precise for small populations and is largely assumption-free but it does not scale up well with population size unless dimensionality reduction methods are used with it. However, neuroscience literature has consistently shown the power and value of considering pairwise or small-group interactions between larger networks to get insights into whole networks [32, 13]. We support the feasibility of this approach simulating discovery of interactions from pairwise-source to single-neuron targets within a 6-neuron network using bias corrections (Fig. S13). We show that using out bias corrections allows discovering the true pairs of neurons that transmit synergistically information even with small numbers of trials, whereas PID without bias corrections would find widespread artificial synergy due to bias.

## 8 Acknowledgements

We are grateful to Christopher Harvey, Patrick Kanold and their laboratories for collaborative work on neural population coding published in previous papers that motivated some of the research questions investigated here. This work was supported by the NIH Brain Initiative grant U19 NS107464 (to SP and TF), the NIH Brain Initiative grant R01 NS109961 and R01 NS108410 (to SP), the Simons Foundation for Autism Research Initiative (SFARI) grant 982347 (to SP), the European Union’s European Research Council grants NEUROPATTERNS 647725 (to TF) and cICMs ERC-2022-AdG-101097402 (to AKE). The Funders had no role in study design, data collection and analysis, decision to publish, or preparation of the manuscript. Views and opinions expressed in this article are those of the authors only and do not necessarily reflect those of the Funders. None of the Funders or granting authorities can be held responsible for them.

## 9 Author contribution

LK and GML contributed equally. SP conceived and supervised the project. LK, GML, NME wrote and developed the algorithms and software. LK, GML, NME, SP performed computations and analyses. All authors contributed materials and methods. SC, SBM and TF contributed and curated data. AKE, TF and SP provided funding and research infrastructure. LK, GML, NME, MC and SP wrote the paper and interpreted results. All authors revised and commented on the text.

## SM1 Appendix / supplemental material

### SM1.1 Definitions of Shannon Information quantities and further relationships with PID components

As described in the main text, once a definition of any of the four PID components is provided, the other three can be obtained as linear combinations between Shannon Information quantities and the defined PID component. To facilitate implementation by users, in the following we report the equations to compute *RI* and *UI* as the linear combination of Shannon Information quantities and *SI* (for which we provided a definition in the main text Eq. (7)).

By subtracting Eq. (4) from Eq. (3) we obtain:

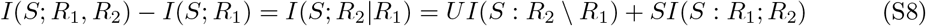

where *I*(*S* : *R*_2_|*R*_1_) is the conditional mutual information about *S* provided by *R*_2_ given *R*_1_, defined as the joint information carried by *R*_1_ and *R*_2_ minus the information carried individually by *R*_1_ [26]. Analogously, by subtracting Eq. (5) from Eq. (3) we obtain:

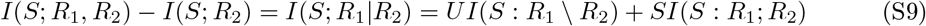

Finally, by subtracting Eq. (4, 5) from Eq. (3) we obtain:

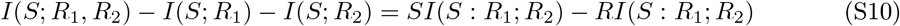

where the quantity on the LHS of Eq. (S10) is known as the co-information *CoI*(*S*; *R*_1_; *R*_2_) of *S, R*_1_ and *R*_2_:

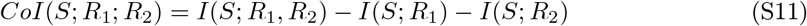

Linear relationships between the four PID components and Shannon information quantities are depicted in Fig. S1. While Eq. (S8-S10) do not impose any independent constraint on the four PID components additionally to the ones in Eq. (3-5), they express explicitly *RI* and *UI* as combinations of Shannon information quantities and *SI*, as follows:

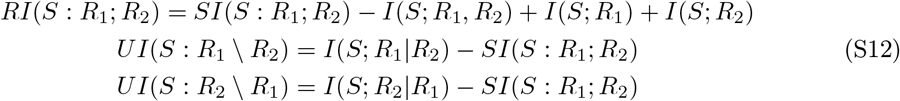

Expressions similar to the ones of Eq. (S12) can be easily obtained to write any three of the PID components as an explicit function of the fourth one. For example, we could have written 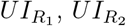, and *SI* as a function of *RI*.

### SM1.2 Definitions and properties of different PIDs

In this section we first provide a mathematical definition of the three measures of PID components used in this paper. Then, we detail important properties of PID of two source variables that are relevant to this study, and discuss which properties these three measures do or do not satisfy. For extensive reviews on available PID measures and their properties, we refer [27, 28].

**Figure S1:**
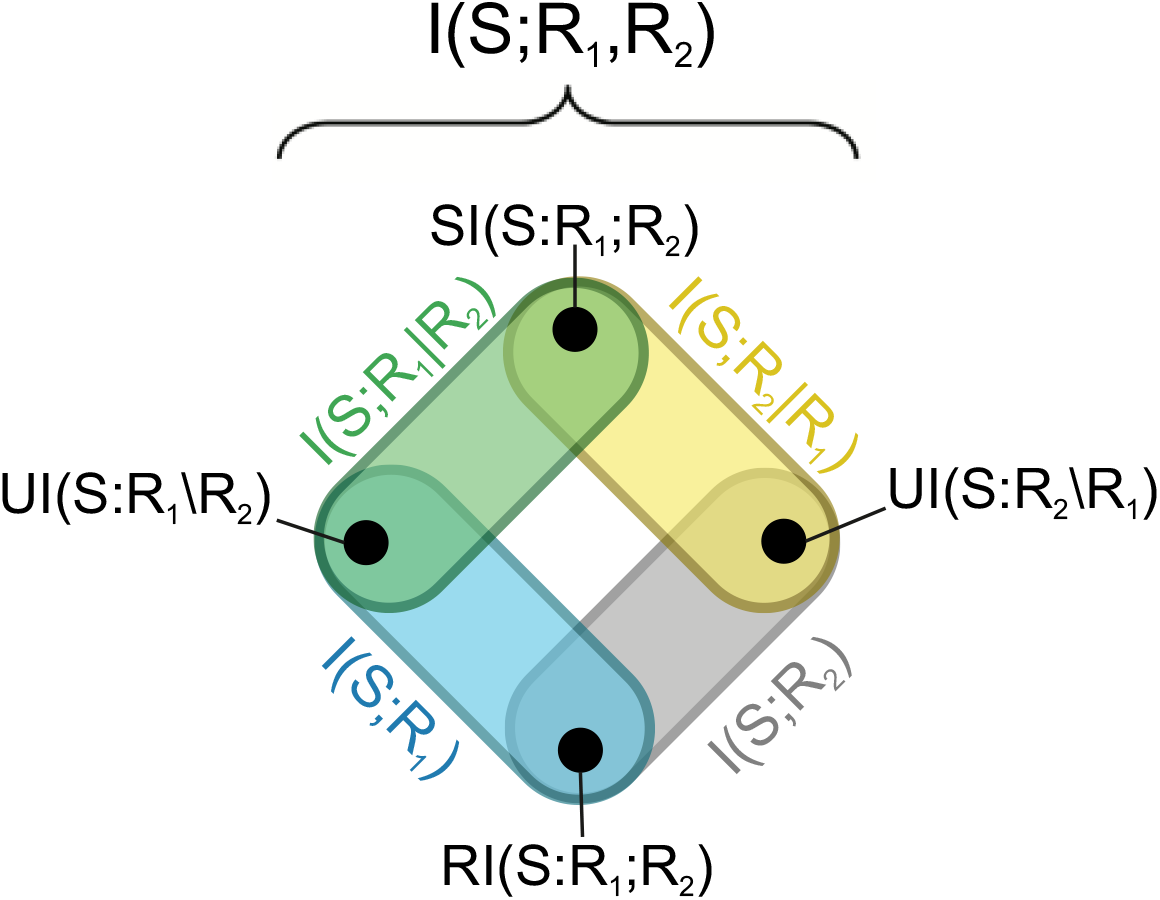
PID decomposes the joint information that the sources *R*_1_ and *R*_2_ carry about *S* into four components: the redundancy *RI*(*S* : *R*_1_; *R*_2_), the unique information *UI*(*S* : *R*_1_ \ *R*_2_) and *UI*(*S* : *R*_2_ \ *R*_1_), and the synergy *SI*(*S* : *R*_1_; *R*_2_). The colored outlines represent the four linear Eq. (4-5) and (S8-S9) that relate the four PID components to Shannon information quantities.

#### SM1.2.1 Different measures of PID components and their properties

***I***_***min***_ In the work introducing PID, Williams and Beer proposed a measure of redundant information about *S*, called *I*_*min*_, that for two source variables *R*_1_ and *R*_2_ is defined as follows:

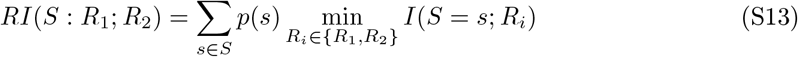

where *I*(*S* = *s*; *R*_*i*_) is the specific information that source *R*_*i*_ carries about a specific value of the target variable *s* ∈ *S*, and is defined as:

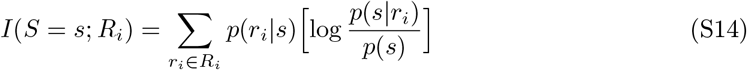

Intuitively, the redundant information computed as in Eq. (S13) quantifies redundancy as the similarity between *R*_1_ and *R*_2_ in discriminating individual values of the target *S*.

***I***_***MMI***_ Minimum mutual information *I*_*MMI*_ was introduced in [21]. This measure is important since both *I*_*min*_ and BROJA measures reduce to *I*_*MMI*_ for Gaussian systems with an univariate target. This is probably the simplest PID measure, defining the redundant information that *R*_1_ and *R*_2_ carry about *S* as follows:

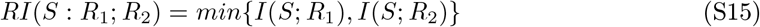

*I*_*MMI*_ quantifies redundant information by directly comparing the amount of information encoded individually by each source, and results in at least one component of unique information always being null.

##### BROJA

The BROJA measure [20] is defined as an optimization problem, minimizing Shannon information quantities that depend on the probability distributions *p*(*S, R*_1_, *R*_2_) defined in the probability space Δ_*P*_ of distributions *q*(*S, R*_1_, *R*_2_) having pairwise marginals between each source and the target equal to the original ones, i.e. *q*(*S, R*_1_) = *p*(*S, R*_1_) and *q*(*S, R*_2_) = *p*(*S, R*_2_). The rationale of this approach is that the synergy can be conceptualized as the information about the target in the joint space of the two sources that cannot be possibly recovered by observing one source at a time (thus from the marginal probabilities), thus it can be defined operationally as the difference between the original joint information and the minimal joint information about the target that can be found in the space Δ_*P*_ of distributions that preserve the marginals (which is the union information, Eq. 6):

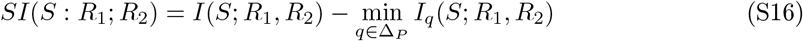

Redundancy and unique information components can then be computed from the synergy and the Shannon information values using the linear constraints as explained above (see Eq. (S12)). Among several advantages of the BROJA PID, we note that this definition insures that the redundancy and the unique information values will have the same value for all the distributions in the space Δ_*P*_ of distributions that preserve the marginals (the so called pairwise marginals property of PIDs).

#### SM1.2.2 Key properties of PID components for two source variables

In their paper introducing PID for the first time, Williams and Beer [12] proposed a set of properties that a measure of *RI* should satisfy. In the following years, other authors introduced new properties which they thought to be important for measures of PID components [19, 29, 28]. Three particularly important properties of PID measures with two source variables are: non-negativity, symmetry and additivity [18].

- Non-negativity: each PID component should be non-negative, i.e. 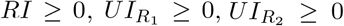 and *SI* ≥ 0. This property is important since it allows interpreting PID components as fractions of the joint mutual information about the target *S* encoded by the source variables (i.e., the average reduction of the entropy of *S* obtained when measuring the two sources simultaneously). All three BROJA, *I*_*min*_ and *I*_*MMI*_ measures satisfy non-negativity [12, 20, 21].
- Symmetry: *RI* and *SI* are symmetric under reordering of source variables, i.e. *RI*(*S* : *R*_1_; *R*_2_) = *RI*(*S* : *R*_2_; *R*_1_) and *SI*(*S* : *R*_1_; *R*_2_) = *SI*(*S* : *R*_2_; *R*_1_). This property is important since also Shannon information quantities which can be decomposed in terms of *RI* and *SI* (including the joint information and the co-information) are symmetric under reordering of *R*_1_ and *R*_2_. All three BROJA, *I*_*min*_ and *I*_*MMI*_ measures satisfy symmetry [12, 20, 21].
- Additivity: given two independent systems of random variables (*S*_1_, *X*_1_, *Y*_1_) and (*S*_2_, *X*_2_, *Y*_2_), then

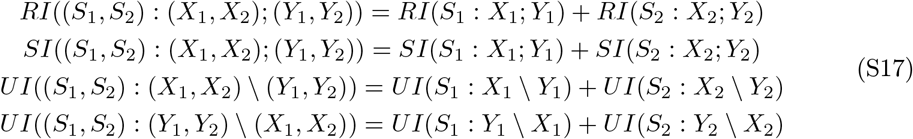

This property is important since it guarantees that PID components of two independent systems can be computed independently and then summed together. BROJA satisfies additivity, while *I*_*min*_ and *I*_*MMI*_ do not [29].

An additional property satisfied by all three measures is the *pairwise marginals* property [28], i.e. *RI*, 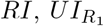 and 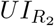 only depend on pairwise marginal distributions between the sources and the target *p*(*S, R*_1_) and *p*(*S, R*_2_). This implies that all dependencies between the target *S* and the sources (*R*_1_, *R*_2_) that go beyond the dependency between the individual sources and the target are quantified as synergy. Even though the pairwise marginals is a fundamental property of the three PID measures, it has been questioned whether PID measures should always satisfy it [28, 67].

### SM1.3 Illustration of bias and details of implementation of the bias correction procedures

#### SM1.3.1 Illustration of the origin of the bias

In this section we illustrate the effect of limited sampling on information calculation (both joint and single neuron information). We simulated two completely stimulus uninformative neurons, responding on each trial with a uniform marginal distribution of spike counts ranging from 1 to 4, regardless of which of 2 stimuli was presented. The neurons are also uncorrelated, so that their joint distribution is uniform across all possible 16 joint responses, and equal between the two stimuli.

Examples of marginal and joint stimulus-conditional neural response empirical response probability histograms sampled from a finite number of trials (50 trials per stimulus in the top row and 200 trials per stimulus in the bottom row respectively) are shown in the left and middle columns (responses to stimuli 1 and 2, respectively). Despite all the true response distributions being uniform across response bins and stimuli, a single instantiation of the probabilities from simulated data will have differences across stimuli that are not real but just due to random fluctuations. With smaller numbers of trials, we can appreciate that the random fluctuations that generate spurious information are larger for the joint probability (the few trials are spread over a larger set of possible responses *R* = 16) than for the marginal probabilities (the few trials are spread over a smaller set of responses *R* = 4). As a result of these spurious differences between stimulus-specific response distributions, the values of information computed from individual simulations with the limited number of trials (right panel) would not be distributed around the true value of zero bits, but around a spurious non-zero value (the bias). The bias is smaller for the single-neuron information values obtained from the marginal probabilities (unique information and single neuron information) than for the information obtained from the joint distribution. With larger number of trials (bottom row), these random fluctuations get smaller (the single instantiation in the bottom row looks more the same across stimuli), but much more so for the marginal than for the joint distribution, and thus the bias reduces more for the quantities obtained from the joint distribution (unique, single neuron) than for the joint information and the synergy.

**Figure S2:**
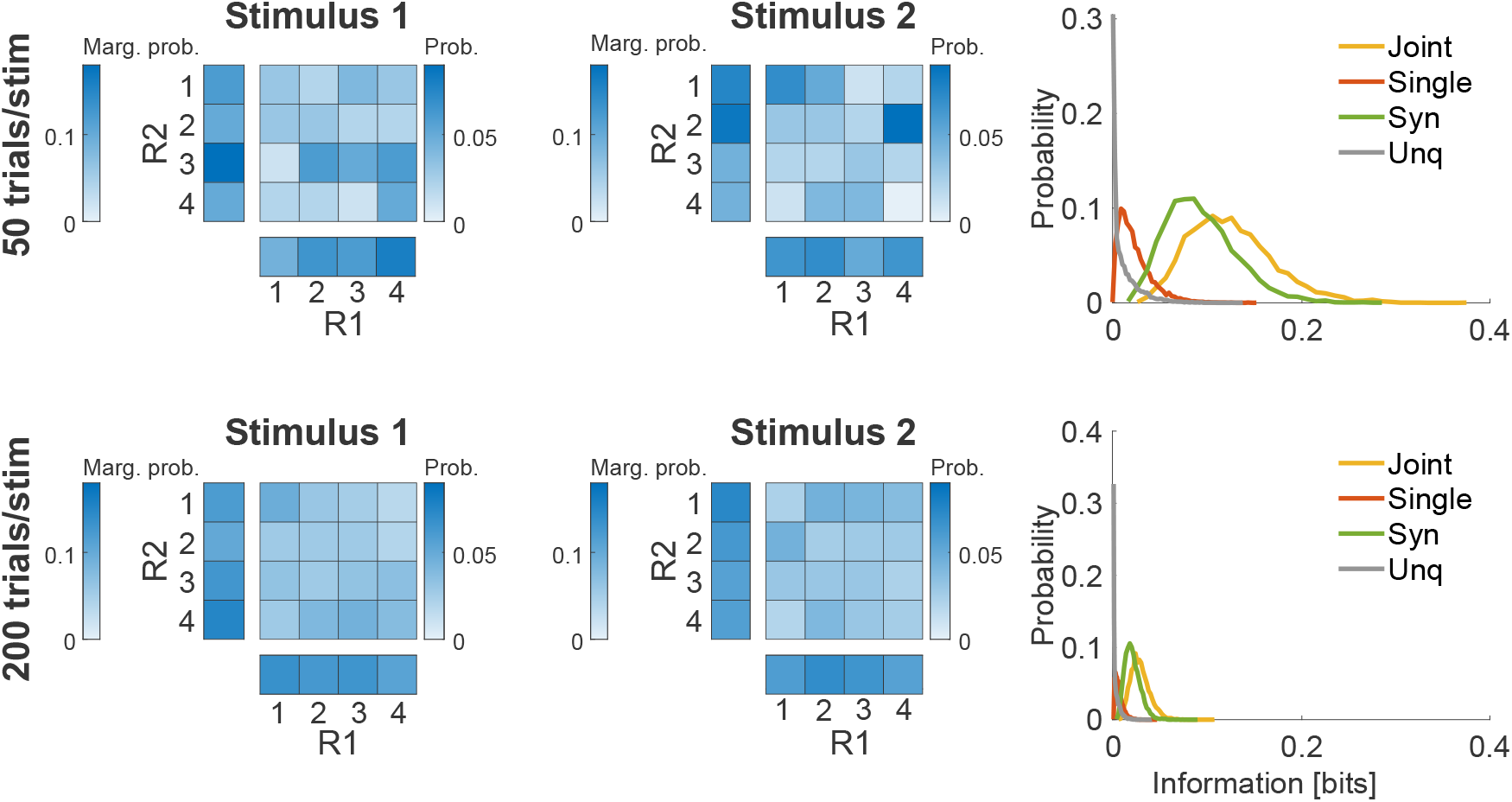
Schematic of the limited sampling bias problem. Two uninformative neurons, responding on each trial with a uniform distribution of spike counts ranging from 1 to 4, regardless of which of 2 stimuli was presented. Examples of empirical response probability heatmaps sampled from 50 and 200 trials per stimulus (top and bottom rows, respectively) are shown in the left and middle columns (responses to stimuli 1 and 2, respectively). Each of the heatmaps has two more heatmaps indicating the marginal probability values. Right: distribution (over 5,000 simulations) of the plugin information values obtained with 50 (top) and 200 (bottom) trials per stimulus respectively.

#### SM1.3.2 Quadratic Extrapolation (QE)

The Quadratic Extrapolation (QE) procedure [43] assumes we are in the asymptotic sampling regime, so that the bias of the entropies and information can be accurately approximated as second order expansions in 1*/N* [14]. That is, we assume that

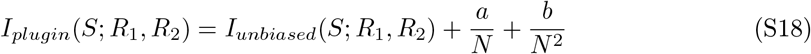

where *a* and *b* are free parameters estimated from the data. This is done by re-computing the information from fractions (*N/*2 and *N/*4) of the data available and then fitting (using a least-square-error procedure) the plugin information values obtained with fractions of data to the preceding quadratic function of 1*/N*. This provides the best-fit estimates of the parameters a and b and consequently the estimate of *I*_*unbiased*_(*S*; *R*_1_, *R*_2_). In the numerical implementation of QE, we allowed for a user-selected number *K* of data partitions into halves and quarters. We performed a separate extrapolation for each partition and then we averaged the resulting estimates of *I*_*unbiased*_(*S*; *R*_1_, *R*_2_) to obtain our final estimate of *I*_*unbiased*_(*S*; *R*_1_, *R*_2_). The procedure is explained in the above for the joint information but we applied it to each and every PID term.

#### SM1.3.3 Shuffling-subtraction bias correction

To estimate the level of the bias, we computed PID after randomly permuting the stimulus values across trials. In this way, all information about the stimulus is lost and all PID terms should be zero, but for finite sampling effect. Thus, the resulting PID terms can be taken as an estimation of their bias, which we subtracted from their plugin estimates. In our implementation, we allowed the user to perform a number *V* of random permutations and then we took as a measure of the bias (to be subtracted from the plugin estimate to obtain the unbiased estimates) the average of the PID term computed over all random realizations of the permutation.

In simulations in the main text and supplement, we used *V* = 20 and *K* = 20. In the analyses of real data, we used *V* = 1 and *K* = 1.

### SM1.4 Details of the binning procedures used to compute the PID

For all simulated data, we estimated the PID components based on the following binning approach. First, in each simulation, we generated a large number (*n* = 2048) of trials per stimulus for each set of model parameters. Then, we discretized the probability distribution of spike counts for each neuron in the chosen number of bins. We computed the bin edges that made the partition of trials as equally populated as possible, following the procedure of Ref. [68]. These bin edges were then used for all simulations computing PID as a function of the number of trials per stimulus. We chose to fix the bin edges for each simulation to ensure that differences across trial numbers in the estimated values of the PID components were due only to trial numerosity and not to other reasons. This was important to study the properties of the information estimates as a function of the number of available trials.

For real data, the binning procedure is specified in the SM section describing their information analysis (see SM Section SM1.10).

A table with all the number of bins used to discretize single neuron activity in each figure is reported in the Table below.

**Table S1:** Number of bins used to discretize neural activity for information-theoretic analyses of simulated and real data, for both the main text and SM figure.

| Number of bins | 2 | 3 | 4 | 8 |
| --- | --- | --- | --- | --- |
| **Figures** | Fig.4C<br>Fig.S4 | Fig.4A<br>Fig.4B | Fig.1<br>Fig.2<br>Fig.3<br>Fig.S2<br>Fig.S3<br>Fig.S6<br>Fig.S7<br>Fig.S8<br>Fig.S9<br>Fig.S10<br>Fig.S11<br>Fig.S12<br>Fig.S13 | Fig.S5 |

### SM1.5 Details of simulations used to test the bias properties and further results of these simulations

#### SM1.5.1 Details of simulations

To test the bias of the PID algorithms, we developed a simulation of the spiking activity of two neurons responding to a set of stimuli, as follows. The spike count *r*_1_ and *r*_2_ of each of the two simulated neurons for each of the four stimulus values was the sum of two Poisson processes, one Poisson process that was independently drawn for each neuron expressing the variability of responses that is “private” to each neuron, and a second Poisson process that is shared between the two simulated neurons and which gives rise to noise correlations.

The equations for the mean rates (that is, mean spike counts) of each Poisson process were as follows:

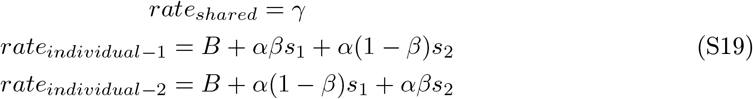

In the above mean rate equation used for our simulations of the spike counts of pairs of neurons in response to the stimulus, we had the following free parameters, which were varied across simulations. We had (i) a parameter *α* regulating the strength of stimulus tuning of the spike counts of the individual neurons or in other words the separation in response strength between least and most effective stimuli (higher values of *α* leading to higher values of individual neuron information and thus to higher values of joint information); (ii) a baseline parameter *B* expressing the overall baseline level of activity common to all stimuli (higher values of *B* in general decreased information of individual neurons and thus the joint information because they increased the standard deviation of responses at fixed stimulus and thus made the rate separation *α* between most and least effective stimuli smaller in standard deviation units); (iii) a parameter *β* which increased the amount of redundancy because it regulated the dissimilarity of tuning of the spike count of each individual neurons to the two stimulus features (*β* close to zero means independent tuning, i.e. each neuron encodes a different feature; higher values of *β* close to *β* = 0.5 indicate that both neurons encode the features similarly and redundantly, i.e. the two neurons encode the same linear combination of features); (iv) and a parameter *γ* regulating the strength of the shared process and thus the strength of correlated firing (increasing *γ* increased synergy because it made the two neurons more strongly correlated, and the fact that the proportion of shared spikes differed between stimuli that were more or less effective for each individual neuron made the overall correlation strength stimulus dependent thereby increasing the amount of synergistic information about the stimulus that can be gained only measuring the joint responses of the two neurons [11]).

For the three scenario presented in the top, middle and bottom rows of Fig. (1, 2, S3, S4, S5, S6, S7, S8, S10, S11) the simulations parameters were as follows. For all scenarios, *α* was set to 7 for the lower information case and to 10 for the higher information case, and *B* was set to 5. For the uncorrelated scenario (top row), we set *γ* = 0 and *β* = 0. For the high redundancy scenario (middle rows), we set *γ* = 2 and *β* = 0.4. For the low redundancy scenario (bottom rows), we set *γ* = 20 and *β* = 0.1. For the figures with the parameter sweeps (Fig 3, S9), we used the same *B, α, β* values as above for the three scenarios but we varied *α* from 1 to 15 in steps of 1.

#### SM1.5.2 Further results of simulations

Here we present and collect all additional analyses of simulated data reporting the bias properties of the PID.

**Figure S3:**
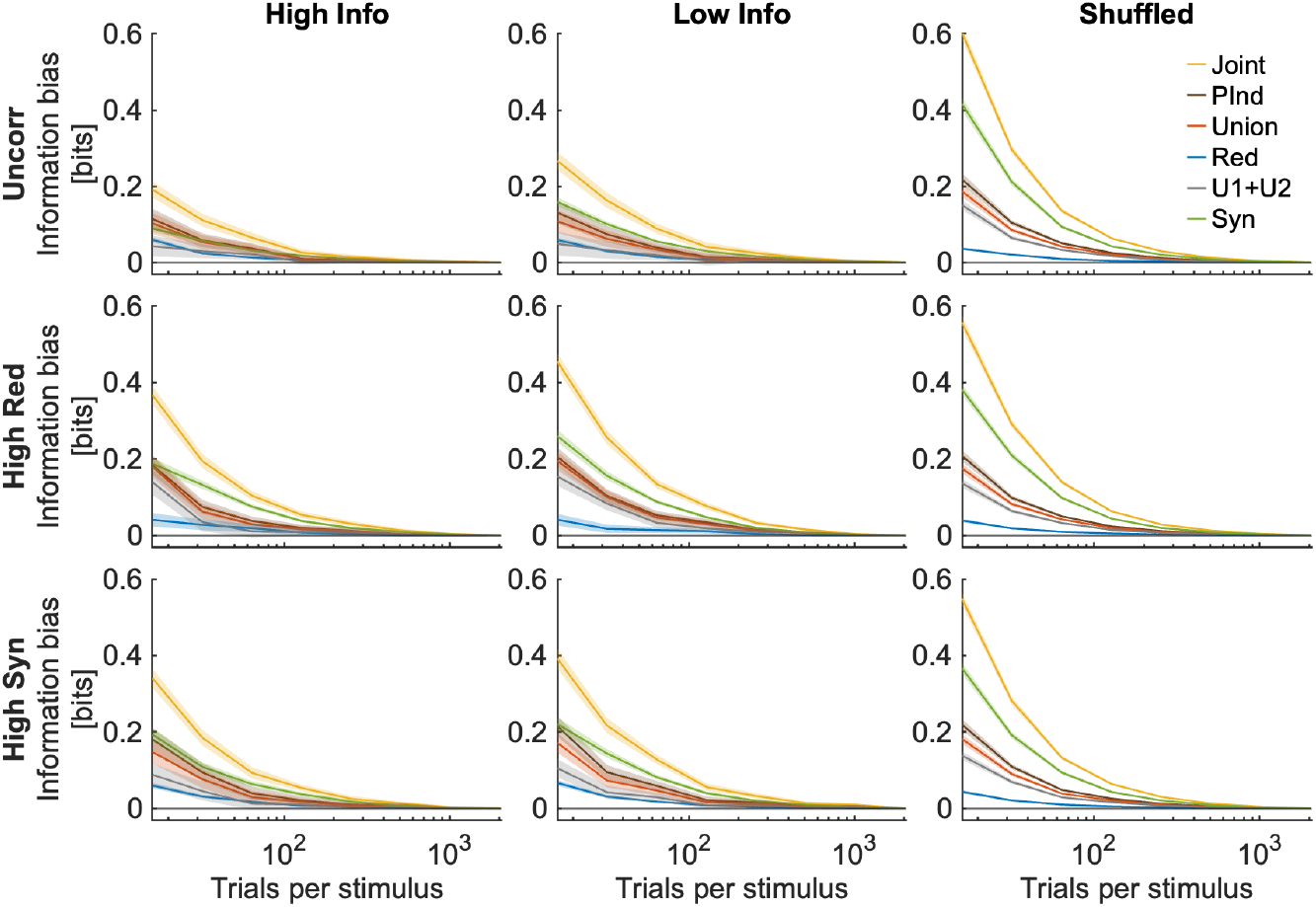
Joint information and PID quantities as a function of the number of simulated trials used to compute them. Here we used the plugin method. In each panel we plot (rather than the information value itself) the information component bias, computed as the information component estimated with the considered number of simulated trials minus the asymptotic information component estimated with the largest number of simulated trials, that is 2048 trials per stimulus). Top, central and bottom row plot the simulated scenarios with no interaction, high redundancy and high synergy, respectively (see SM Section SM1.5). Left, center and right columns represent simulations with higher information (*α* = 10), lower information (*α* = 7) and with shuffled low-information data. “Syn”: synergy. “Red”: redundancy. “U1+U2”: sum of the two unique information of each neuron. We used *R* = 4 discretization bins for each neuron (Table S1). Each panel plots mean ± 2 SEM over *n* = 96 simulations.

**Figure S4:**
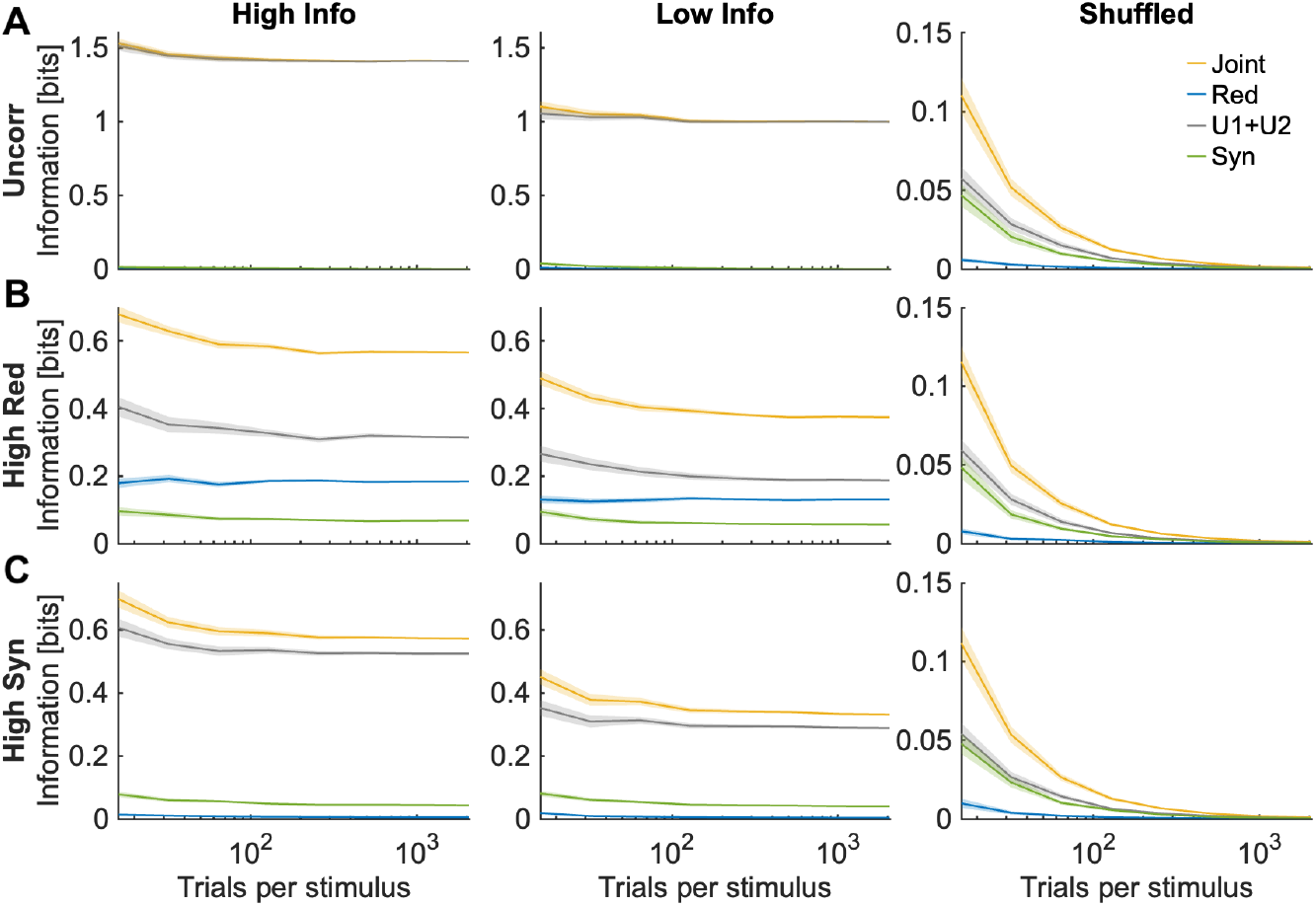
Joint information and PID quantities using BROJA as the redundancy measure as a function of the number of simulated trials used to compute them. Plotting conventions are exactly as in Fig. 1. We used *R* = 2 discretization bins per each neuron (see Table S1). Results in each panel are plotted as mean ± 2 SEM over *n* = 96 simulations.

**Figure S5:**
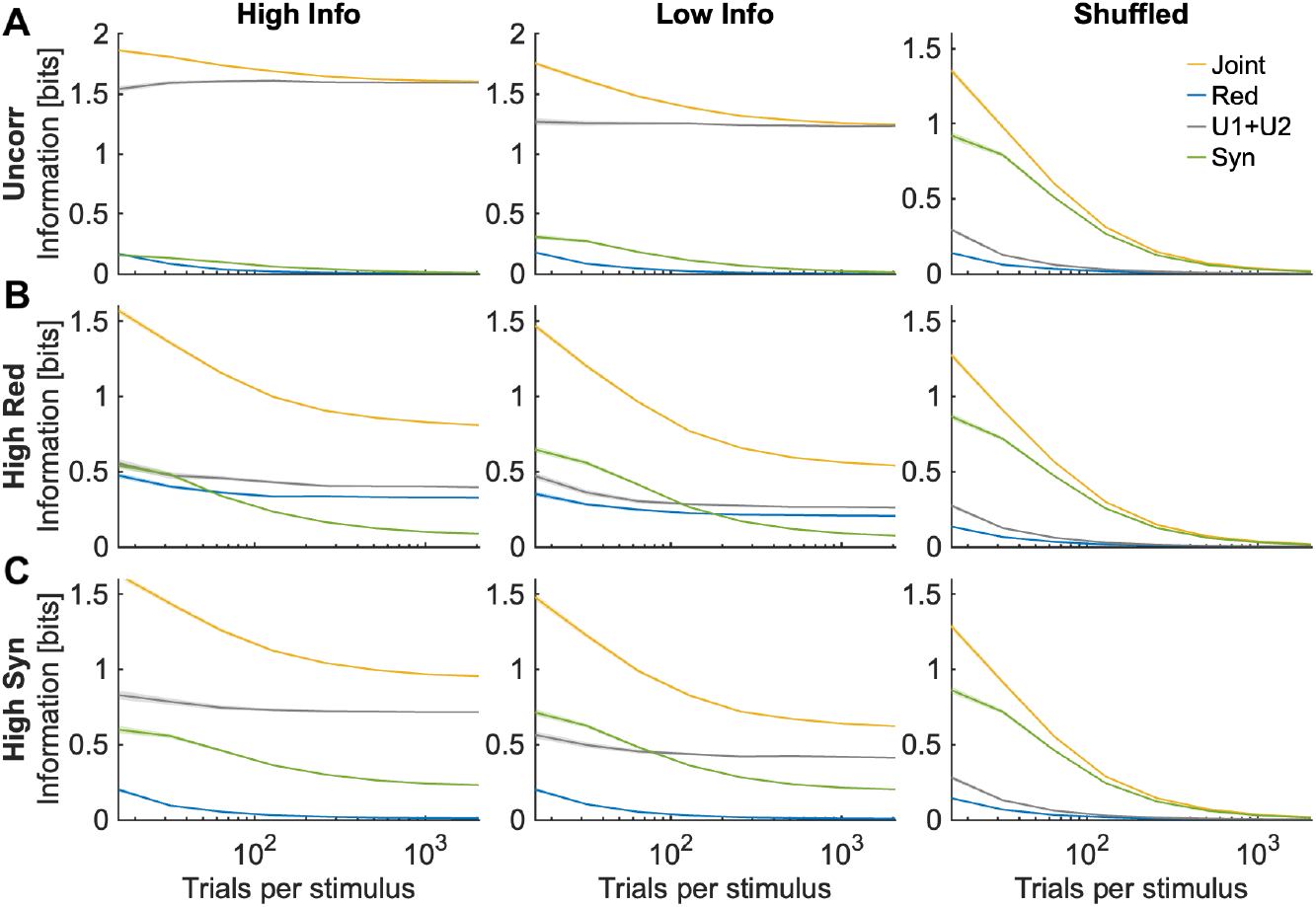
Joint information and PID quantities using BROJA as the redundancy measure as a function of the number of simulated trials used to compute them. Plotting conventions are exactly as in Fig. 1. We used *R* = 8 discretization bins per each neuron (see Table S1). Results in each panel are plotted as mean ± 2 SEM over *n* = 96 simulations.

**Figure S6:**
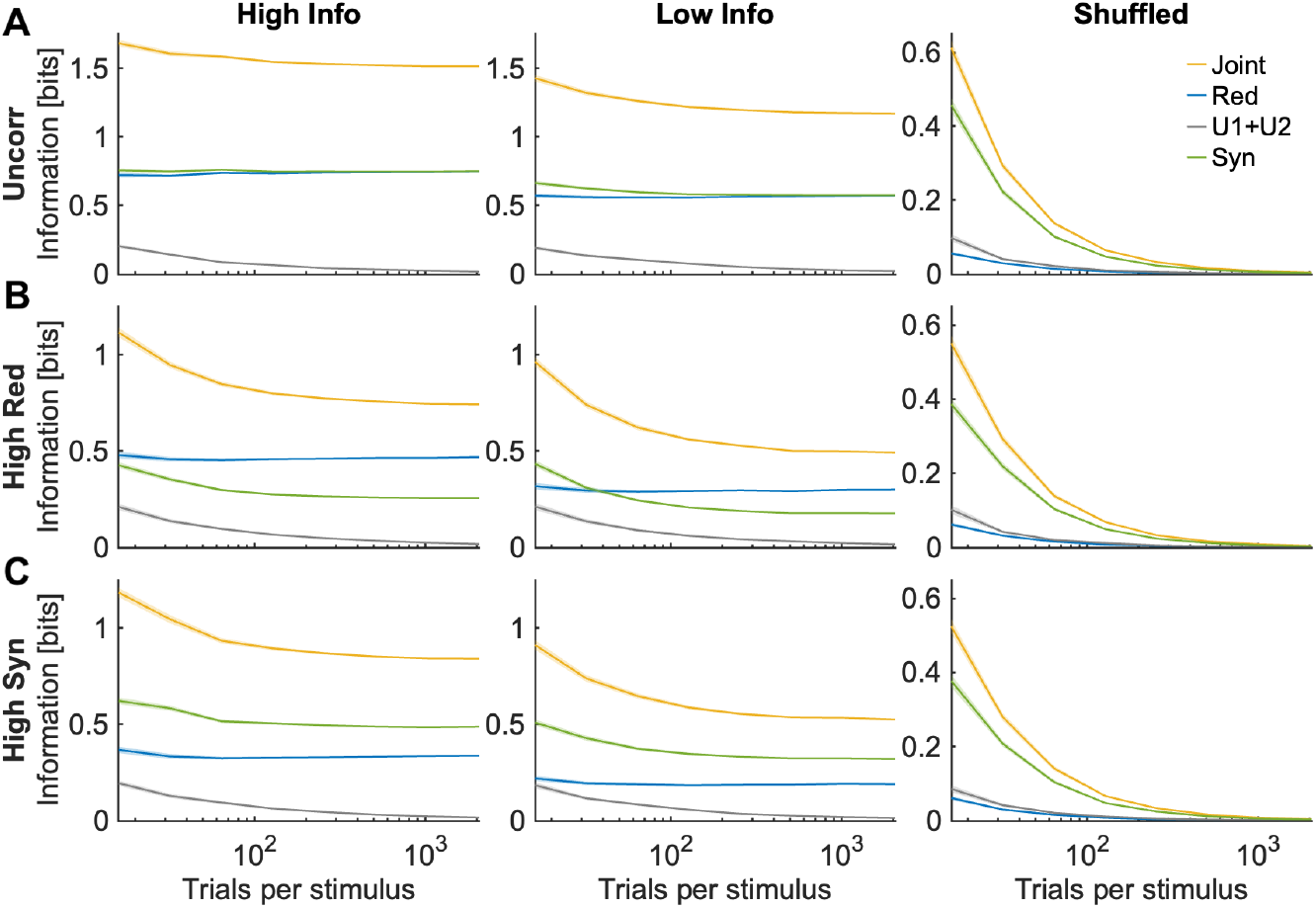
Joint information and PID quantities using *I*_*min*_ as the redundancy measure as a function of the number of simulated trials used to compute them. Plotting conventions are exactly as in Fig. 1. We used *R* = 4 discretization bins per each neuron (see Table S1). Results in each panel are plotted as mean ± 2 SEM over *n* = 96 simulations.

**Figure S7:**
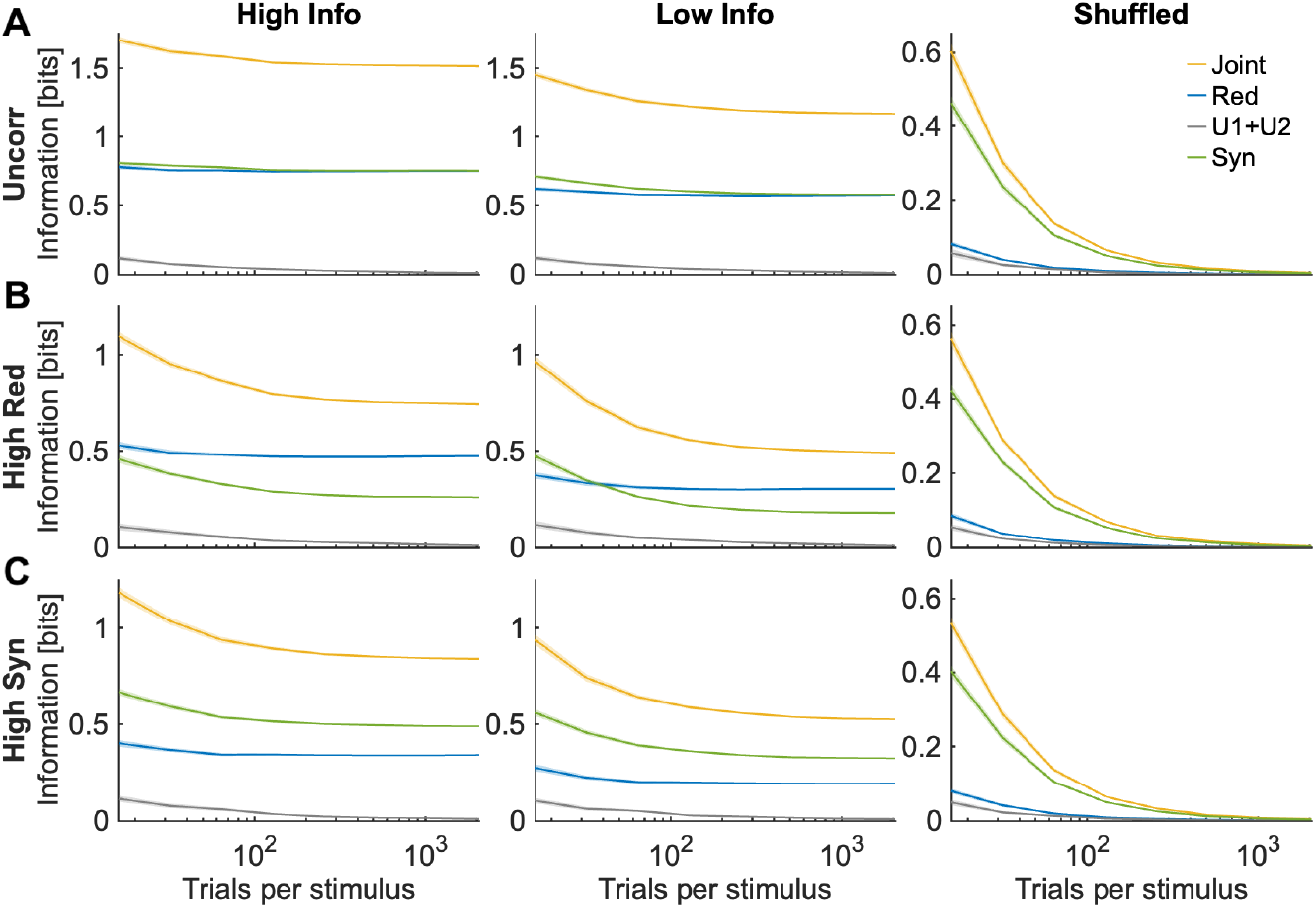
Joint information and PID quantities using *I*_*MMI*_ as the redundancy measure as a function of the number of simulated trials used to compute them. Plotting conventions are exactly as in Fig. 1. We used *R* = 4 discretization bins per each neuron (see Table S1). Results in each panel are plotted as mean ± 2 SEM over *n* = 96 simulations.

### SM1.6 Further Results Bias Corrections

In this subsection we collate figures reporting additional results of the PID bias correction methods on simulated data.

**Figure S8:**
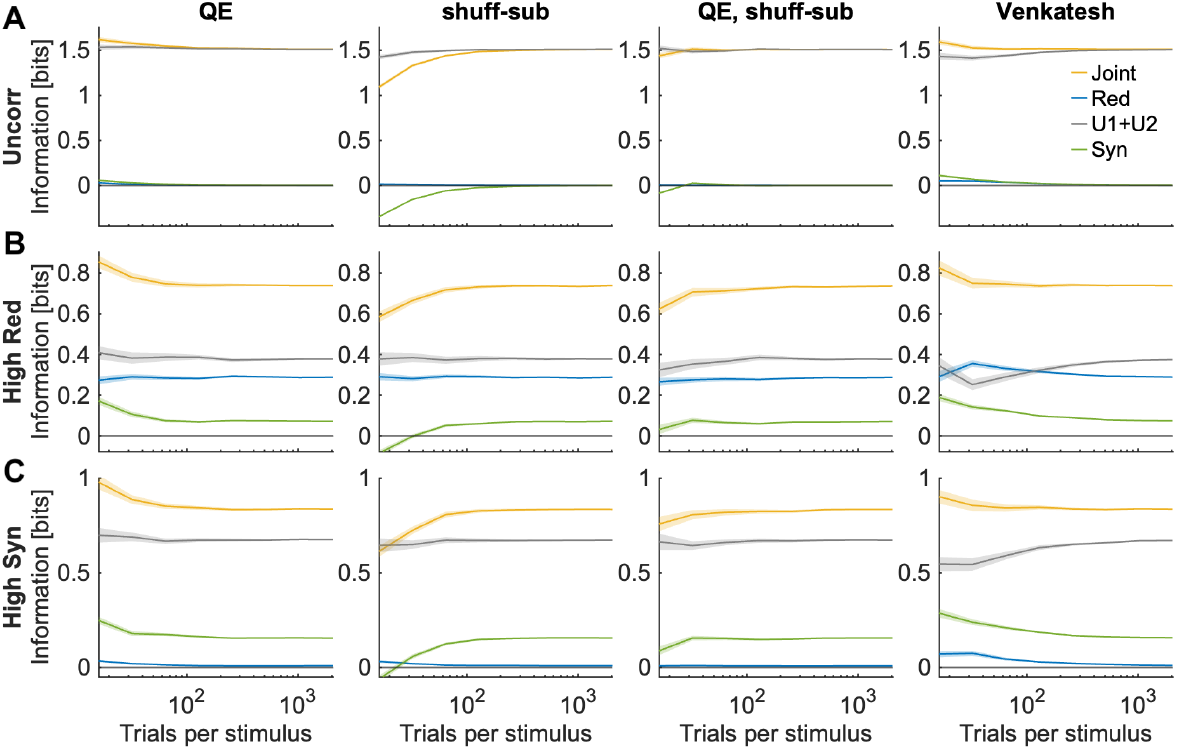
Bias corrections for the case of high information. Joint information and PID quantities as a function of the number of simulated trials used to compute them. Plotting conventions are exactly as in Fig. 2. We used 4 discretization bins for each neuron (see Table S1). Results in each panel are plotted as mean ± 2 SEM over *n* = 96 simulations.

**Figure S9:**
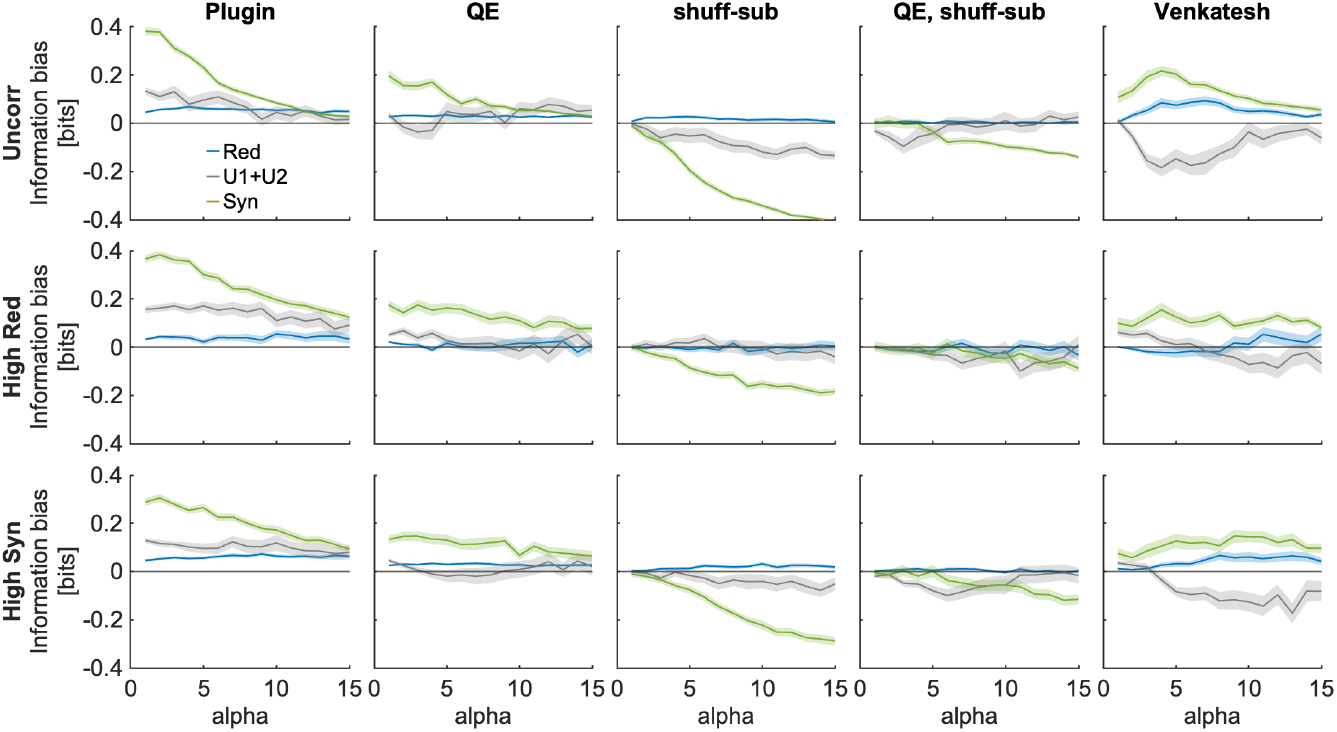
Performance of bias corrections using *N*_*s*_ = 16 trials per stimulus with 4 discretization bins for each neuron, as function of the parameter *α* increasing single-neuron information in the simulated data. In each panel we plot (rather than the information value itself) the information component bias, computed as the information component estimated with the considered number of simulated trials minus the asymptotic information component estimated with the largest available number of simulated trials, that is 2048 trials per stimulus). Plotting conventions are exactly as in Fig. 3.

**Figure S10:**
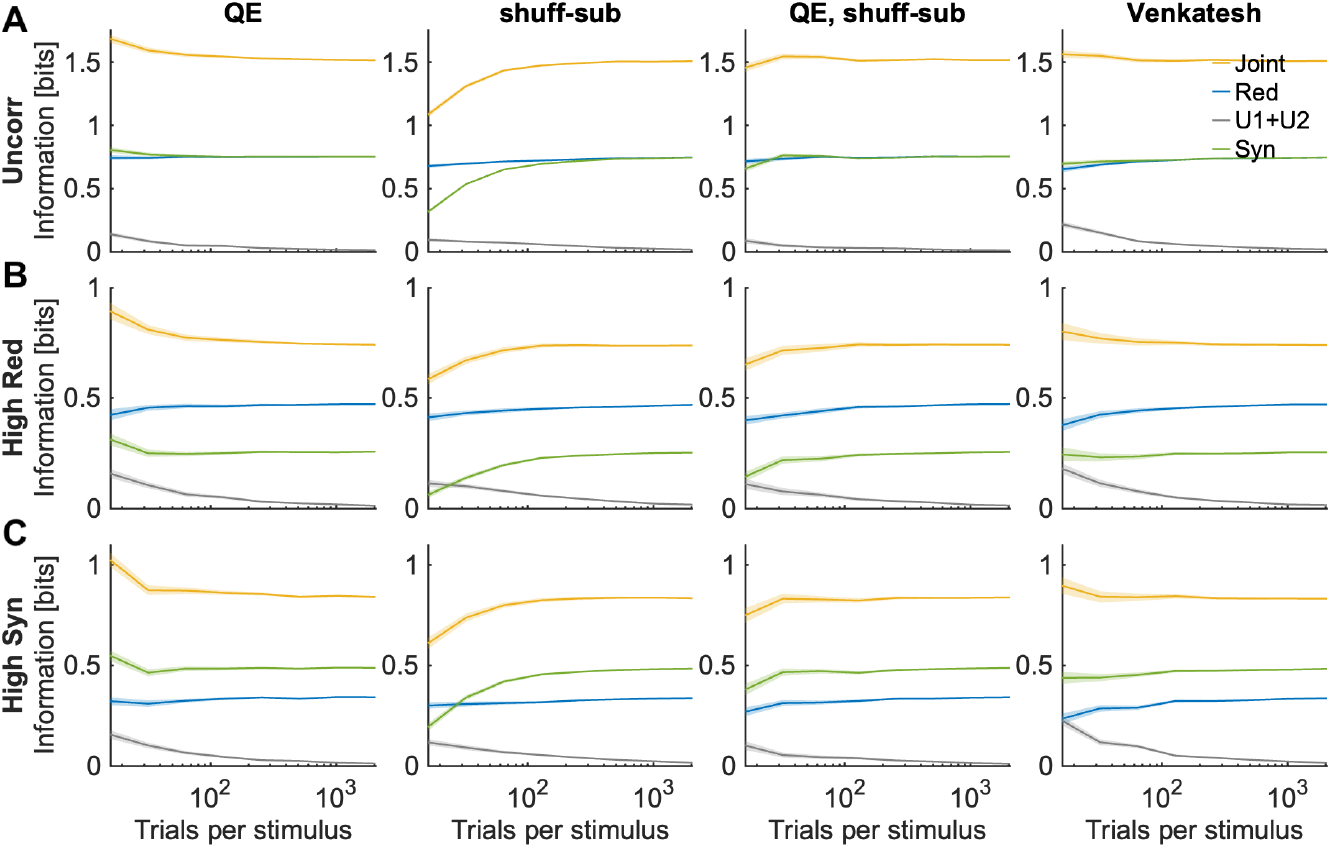
Bias corrections for the case of high information using *I*_*min*_ as the redundancy measure. Joint information and PID quantities as a function of the number of simulated trials used to compute them. Plotting conventions are exactly as in Fig. 2. We used *R* = 4 discretization bins per each neuron (see Table S1). Results in each panel are plotted as mean ± 2 SEM over *n* = 96 simulations.

**Figure S11:**
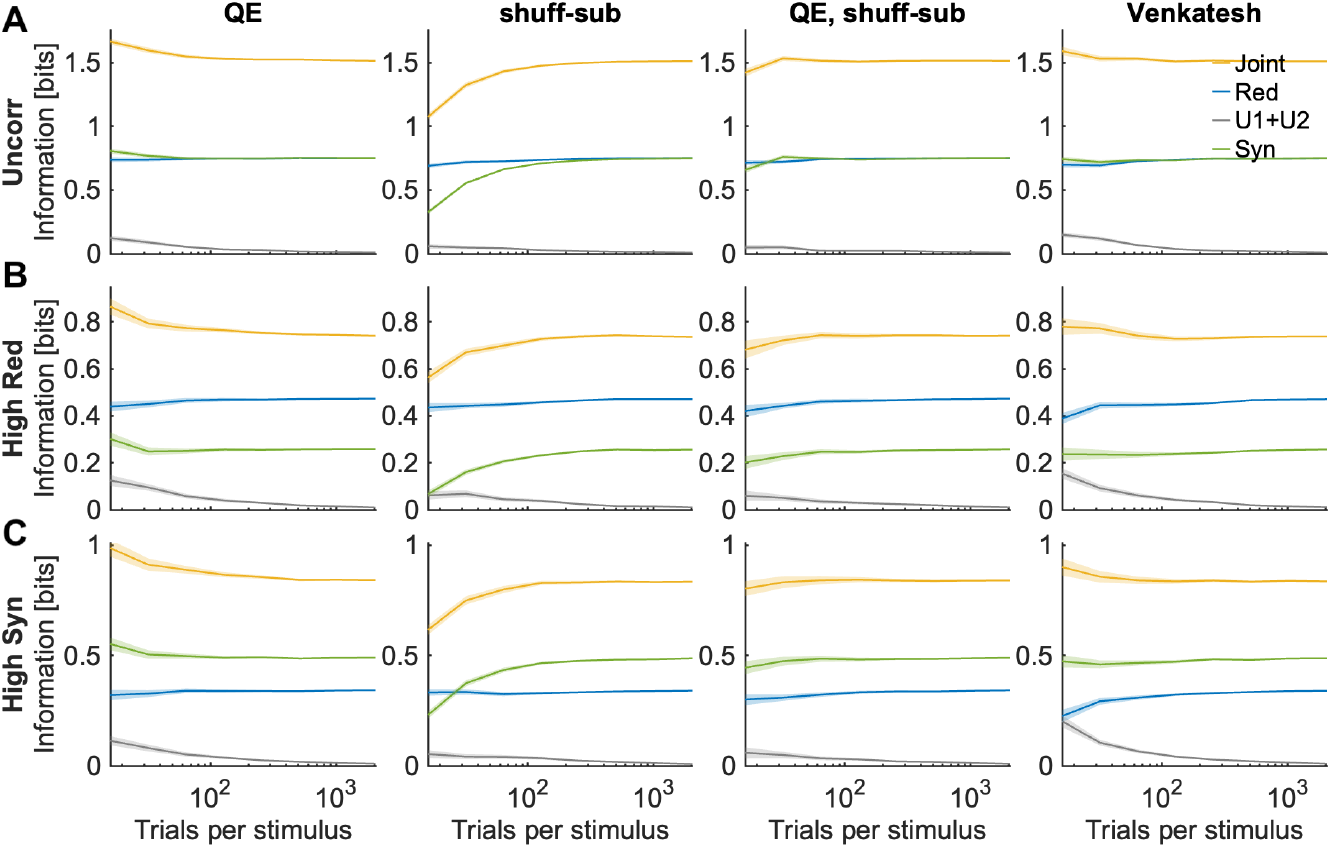
Bias corrections for the case of high information using *I*_*MMI*_ as the redundancy measure. Joint information and PID quantities as a function of the number of simulated trials used to compute them. Plotting conventions are exactly as in Fig. 2. We used *R* = 4 discretization bins per each neuron (see Table S1). Results in each panel are plotted as mean ± 2 SEM over *n* = 96 simulations.

### SM1.7 Simulation showing that the Gaussian approximation for Shannon information fails for neural spike rates with realistically low numbers of spikes

In this subsection, we use simulations of neural activity to exemplify why the Gaussian approximations for information, often used in PID, are not suitable for computing information about stimuli carried by neurons.

We consider here the mutual information about a stimulus *s* carried by the response of one neuron *r*_*i*_, which can be quantified as the difference between the entropy *H*(*R*_*i*_) of the distribution *p*(*r*_*i*_) of responses *r*_*i*_ across of stimuli, called response entropy in the neural literature, and the conditional entropy *H*(*R*_*i*_|*S*) of the stimulus-specific distributions *P* (*r*_*i*_|*s*) referred to as noise entropy in the neural literature:

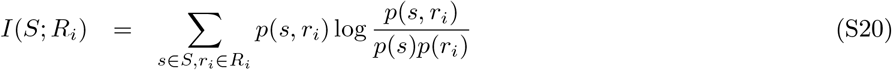

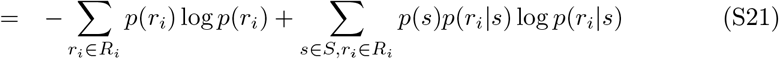

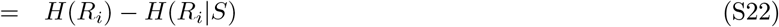

It is well documented that the number of spikes emitted by a neuron in response to different stimuli follows approximately a Poisson distribution [34]. Moreover, the time windows in which neurons process and transmit information are usually short of the order of tens to hundreds of milliseconds [45]. Given that the firing rate of cortical neurons ranges from one or few spikes per second (when considering the less effective stimuli) to several tens of spikes per second (when considering the most effective stimuli), firing rate distributions relevant for neural information processing can be conceptualized as near-Poisson with relatively small average mean spike counts. Under these conditions, probability of neural responses conditional to the stimuli are not well approximated by Gaussian distributions, as often assumed by implementations of PID [21, 69, 32, 18]. As a result, the noise entropy will be overestimated (remember that the Gaussian distribution is the one with the highest entropy across all distributions with a given variance). If the different stimuli presented are few in number (that is, *S* is small) the probability *p*(*r*) of response *r* across all stimuli will be very far from a Gaussian, and as a consequence the response entropy will be overestimated even more than the noise entropy, and as a result the information will be overestimated substantially. Hence, Gaussian PIDs cannot be used to estimate information reliably in typical experiments involving the recording of neural activity, and estimation methods that respect the discrete nature of neural activity will be better suited.

To illustrate this, we simulated spike count responses of an individual neuron in response to a set of *S* = 4 different stimuli defined by two independently-drawn binary features *s*1, *s*2. We simulated stimulus-specific neural spike counts in response to a stimulus defined by the two feature values (*s*1, *s*2) as a Poisson process with average count *r*, as follows:

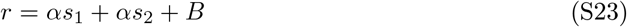

The parameter *B* represents the baseline firing and the parameter *α* represent the separation of the mean responses between the least effective and most effective stimuli. The parameter *α* increases the information (as it increases the separation between least and more effective stimuli), whereas the parameter *B* decreases the information (it increases of standard deviations of stimulus-specific responses, which in turn decreases the separation between responses to different stimuli in units of these standard deviations). We used three approaches to compute information. In the first, we computed the ground-truth information by computing the probabilities of each single possible spike count value, which captures perfectly the information expressed by the above Poisson process; the information computed by discretizing the responses into *R* = 4 equipopulated bins (at it could be done reliably also in experiments with more limited numbers of trials) and using the Gaussian approximation (the latter is computed using the information that would be obtained if all probabilities *P* (*r*|*s*) and *P* (*r*) were Gaussian with the standard deviation *σ* and *σ*_*s*_ of measured empirically from the empirical *P* (*r*_*i*_) and *P* (*r*_*i*_|*s*), as done in Gaussian PID [69, 32, 18]:

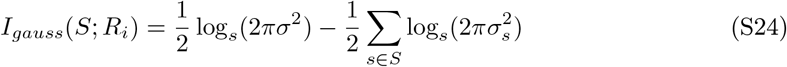

In Fig. S12 we plot the values of information obtained with these 3 approaches simulating neural responses across values of the parameters *α* and *B* (*α* and *B* were varied between 2 and 14 in steps of 1). We computed information using a large number (10000) of simulated trials per stimulus to avoid bias issues. The Gaussian information presented a large overestimation of information with respect to the ground truth information encoded by the Poisson process. On average across all simulated cases, the values of Gaussian information overestimated the ground truth values by 21%.

In contrast, a simple discretization into *R* = 4 equipopulated bins of the neural responses led to a very precise estimation of the ground truth information. On average across all simulated cases, the values of the *R* = 4 discretization underestimated the ground truth values by only 2% (the slight underestimation due to a coarser discretization of discrete data is due to the data processing inequality).

### SM1.8 Asymptotic expansion of the bias of the PID components in the limit of large numbers of experimental trials

To derive analytical approximations to the bias, we make the assumption of a large *N* limit (where *N* is the total number of experimental trials available to compute the probabilities), defined formally as the case when *N*_*s*_*P* (*r*_1_, *r*_2_|*s*) ≫ 1 for each stimulus *s* and joint response *r*_1_, *r*_2_, where *N*_*s*_ is the number of trials for each stimulus used to compute the stimulus-specific neural response probabilities. (If this is satisfied then it is also satisfied that *NP* (*r*_1_, *r*_2_) ≫ 1 for each joint response *r*_1_, *r*_2_, and that the same applies to all the marginal single-neuron probabilities, e.g. *N*_*s*_*P* (*r*_1_|*s*) ≫ 1). While it is unrealistic that this large *N* limit case will be formally satisfied in practical experiments, this type of derivation has proven very useful to understand the properties of the bias and to design strategies to cope with it [64, 14, 15].

Suppose we have *M* trials (for us, *M* will be either *N*_*s*_ or *N* depending on whether we will consider noise or response entropy) and we want to compute the entropy in a certain discrete space *X* (for us, this space will be the joint space of pairwise neural activity or the space of marginal distributions of individual neural activity). The entropy of the probability distribution is defined as

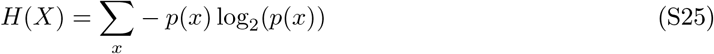

where it is important to bear in mind that the sum over possible values of *x* for the discrete entropy includes only values of *x* with probability larger than zero.

The values of the probabilities are not known a priori in experimental situations. They could be perfectly computed from the data if we had infinite amounts of experimental trials, but we assume that we have only a finite amount of trials *M*. If we compute the probabilities from the empirical occurrences *p*_*M*_ (*x*) from the *M* available trials, we have the value of the entropy *H*_*M*_ obtained with a finite number of trials *M* :

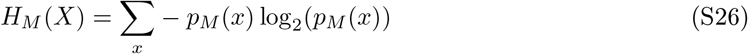

**Figure S12:**
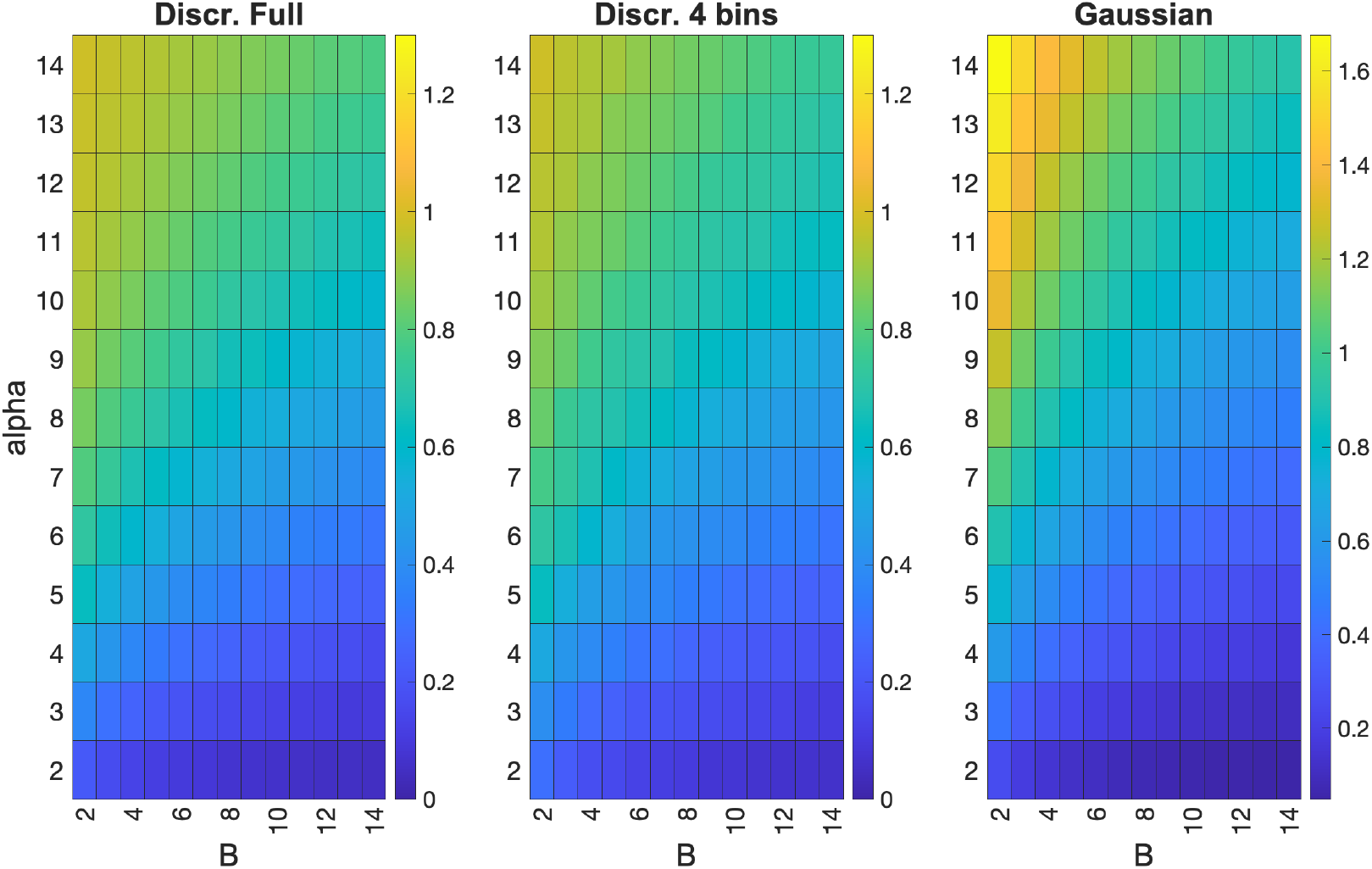
Comparison between discrete and Gaussian computations of the stimulus information carried by an individual Poisson neuron. We report the information values obtained when evaluating the stimulus information carried by a Poisson neuron simulated with mean spike count set Eq. (S23) as a function of the parameters *B* and *α*. We simulated 10000 trials for each of the 4 stimuli. Left: ground truth information of the Poisson neuron obtained with discrete probabilities considering the probabilities of all possible spike count values. Middle: information computed with the discretized approach by binning the possible spike counts into 4 equipopulated bins. Right: information values computed with the Gaussian approximation of the distributions of spike counts. The Gaussian approach strongly overestimates the ground-truth values of information (by 21% on average across all simulated parameters), whereas the discrete approach using the coarse binning is far more precise (within 2% on average across all simulated parameters)

We would like to compute the finite sampling bias of the entropy, defined as the difference between the average value ⟨*H*_*M*_ ⟩_*M*_ of *H*_*M*_ (over many instantiations with *M* trials) and the true value *H* obtained when the true distributions are known. In what follows, we will compute an approximation to the bias in the large *M* limit, following the procedure of [14]. However, other methods lead exactly to the same result for the leading approximation to the bias in the large *M* limit [64, 15]. To compute analytical approximations to the bias of the entropy, we will make use of the logarithm expansion:

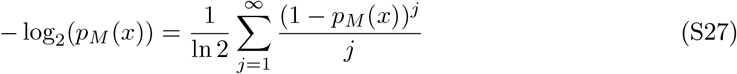

This is convergent for any value of 0 *< p*_*M*_ (*x*) *<* 1 (note that values of *p*_*M*_ (*x*) = 0 provide a vanishing contribution (thus do not enter) the above sum and if *p*_*M*_ (*x*) = 1 for one response then the entropy is trivially zero). This allows to rewrite the entropy computed from *M* samples as a sum over powers of the probabilities:

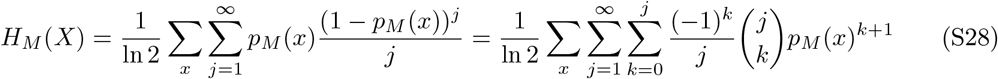

From the above equation, it follows that averages over all instantiations with *M* trials of the entropy computed with *M* trials can be obtained from the averages of the powers of the probabilities over instantiations with *M* trials. Because we use a discrete computation, we assume without loss of generality that the neural response and the stimulus-specific response probabilities follow a multinomial probability with arbitrary parameters. The value of the probability *p*(*x*) computed from the empirical occurrences *p*_*M*_ (*x*) using *M* trials on average overall all possible outcomes is unbiased, as its value ⟨*p*_*M*_ (*x*)⟩_*M*_ coincides with the true value *p*(*x*)that would be estimated with an infinite number of trials:

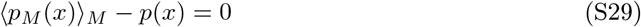

When computing with *M* trials the power *k* (for *k >* 1) of a discrete probability *p*(*x*), on average over all possible outcomes the power of the empirical probability has instead a bias. Under the assumptions that the number of trials *M* is so large that *Mp*(*x*) ≫ 1 for all *x*, the bias of the power *k* of the probability has the following asymptotic expression [70, 14]:

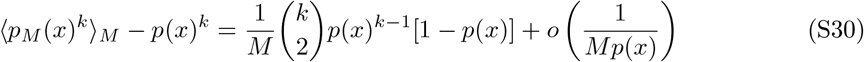

Higher orders terms of the bias of the probability powers can be computed analytically and are available in Ref. [70]. By inserting the above expansion of the average probability with *M* trials into the expansion of the entropy one can obtain by some algebra an expansion of the bias of the entropy in inverse powers of 1*/M*

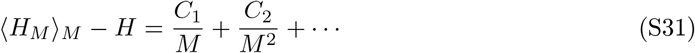

with the leading term 1*/M* in the bias being

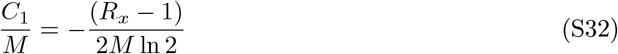

where *R*_*x*_ denotes the number of relevant events (responses) for the considered distribution, defined as the number of responses *x* with probability *p*(*x*) *>* 0. Thus, although the bias is not the same for all probability distributions, in the asymptotic sampling regime, its leading term depends on some remarkably simple details of the response distribution (the number of trials and the number of relevant responses). This makes it possible to derive simple rules of thumb for estimating the bias magnitude and compare the relative bias in different situations. As we will discuss now, this equation can predict very effectively how bias properties of PID terms differ between them within a simulated scenario and how they change across different simulated scenarios.

The leading bias terms of the joint information can be computed by writing it as the difference between joint response entropy and joint noise entropy,

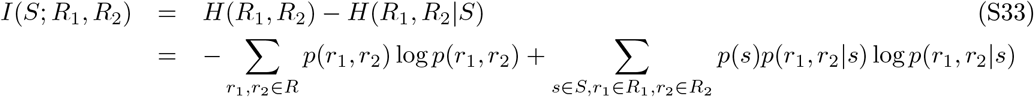

and using the equations above with *M* equal to *N*_*s*_ for the noise entropy and to *N* for the response entropy. In this way we obtain

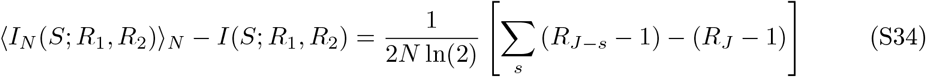

The above bias of *I*(*S*; *R*_1_, *R*_2_) originates from finite sampling fluctuations of the joint probabilities and thus is function of *R*_*J−s*_, the number of relevant bins of the joint stimulus-specific distribution in the joint *r*_1_, *r*_2_ space, *R*_*J*_ is the number of relevant bins of the joint stimulus-unconditional distribution in the joint *r*_1_, *r*_2_ space, and *N* the total number of trials available across all stimuli.

Consider the bias of the single cell entropies (Eq. (S22) above). They depend on the marginal distribution *p*(*r*_*i*_|*s*) and *p*(*r*_*i*_). Thus this bias depends on the finite sampling fluctuations of the marginal probabilities, which will be smaller than those for the joint probability because the available trials are concentrated in a smaller space (Fig. S2). Thus, the bias of the single cells will be much smaller than that of the joint information and its leading term is as follows:

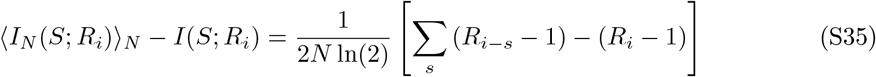

In the above *R*_*i−s*_ and *R*_*i*_ are the number of bins relevant for the marginal stimulus conditional and stimulus-unconditional probabilities. These bin numbers will be less than the corresponding ones for the joint probabilities *R*_*J−s*_ and *R*_*J*_. For example, if we discretized responses into 5 bins per neurons, we expect the joint response bins *R*_*J−s*_ and *R*_*J*_ to be 25 or less, and we expect *R*_*i−s*_ and *R*_*i*_ to be 5 or less. Thus we expect in this case a factor of 5 difference between the joint and individual information bias. In general we can expect the bias of the individual neuron information to be considerably smaller than that of the joint information, with the difference between the two biases getting larger when more bins are used to discretize the activity of each neuron.

Having understood the bias of Shannon information measures relevant for PID, we now use this knowledge to evaluate the bias of the PID terms. We focus first on the bias of the synergy *SI* as defined in BROJA. We remind that the synergy equation is defined as a difference between the joint and the Union information, with the Union information defined as follows:

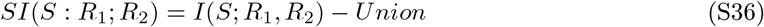

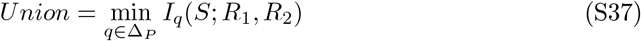

where Δ_*P*_ is the set of all joint probability distributions *q*(*S, R*_1_, *R*_2_) that have the same pairwise marginals *q*(*S, R*_1_) = *p*(*S, R*_1_) and *q*(*S, R*_2_) = *p*(*S, R*_2_) as the original distribution *p*(*S, R*_1_, *R*_2_), and *I*_*q*_(*S*; *R*_1_, *R*_2_) is the joint information computed for distribution *q*(*S*; *R*_1_, *R*_2_). Consider now the Union information. When applied to limited data *M*, the minimization within the Δ_*P*_ space when applied to finite sampling data will tend to select a minimum information within the space set by the marginals *p*_*M*_ (*r*_1_, *s*) and *p*_*M*_ (*r*_2_, *s*) computed with *M* trials. It will thus select probability distributions with a low finite sampling information value. Because this bias is by and large positive, the minimization procedure will tend to select probabilities with lower bias. Among the probabilities with low bias in the Δ_*P*_ space we have the probability *p*_*ind*_(*r*_1_, *r*_2_*s*) defined directly from the marginals such that the neurons would have the same single cell probabilities but no interactions at fixed stimulus (noise correlations)

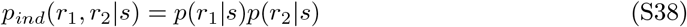

We call the *I*_*q*_ computed on *q* = *p*_*ind*_ the conditionally-independent information *I*_*ind*_(*S*; *R*_1_, *R*_2_), defined as the joint mutual information with stimulus-conditional probabilities *p*_*ind*_(*r*_1_, *r*_2_|*s*) and stimulus unconditional probabilities *p*_*ind*_(*r*_1_, *r*_2_) = Σ_*s*_ *p*_*ind*_(*r*_1_, *r*_2_|*s*)*p*(*s*). (The information is the information carried by the population about the stimulus if the single neuron responses were the same as the original data but there were no noise correlations (no stimulus conditional interactions between the neurons), and it has been used as a reference distribution for understanding whether noise correlations increase or decrease information, see [4, 11].) Importantly for our PID bias understanding, we can use the above bias expansion to compute that in the large *N* limit *I*_*ind*_(*S*; *R*_1_, *R*_2_) has a low bias, very close to the one of the sum of the biases of individual information:

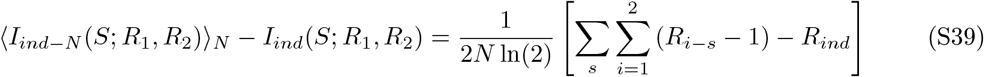

where *R*_*i−s*_ is the number of bins relevant for the marginal stimulus conditional and *R*_*ind*_ is the one relevant for the stimulus-unconditional independent probabilities of the i-th individual neuron (*i* = 1, 2). This will be smaller than the corresponding one for the joint information. For example, if we discretize responses into 5 bins per individual neuron, we would expect 25 bins or less relevant for the joint information and 10 or less (5 for neuron 1 and 5 for neuron 2). In other words, the bias of the joint information grows quadratically with the number of responses of the individual neurons, whereas the bias of *I*_*ind*_(*S*; *R*_1_, *R*_2_) grows linearly. Because the minimal set of finite sampling variations in the Δ_*P*_ space is set by the finite sampling fluctuations of the marginals, we expect and make the assumption that the minimization in the Union calculation will select distributions roughly with the same bias as *I*_*ind*_(*S*; *R*_1_, *R*_2_). To test this, we performed our simulations of pairs of neurons. We found (see Fig. S3) across all conditions and trial numbers that the bias of the Union information is very similar to the bias of *I*_*ind*_(*S*; *R*_1_, *R*_2_). It is actually slightly less which is compatible with the understanding that the minimum selects a similar bias and of course smaller statistical fluctuations.

The synergy is the difference between the joint and the union information. Because the former has a large positive bias (growing quadratically with the number of single-neuron discrete responses) and the latter has a smaller positive bias (growing linearly with the number of single-neuron discrete responses), the synergy is expected to have a relatively large positive bias, smaller than that of the joint information but still large. This prediction is confirmed in all numerical simulations (Fig. 1, 2, S3, S4, S5, S6, S7, S8, S10, S11).

The redundancy is the difference between the sum of single neuron information and the Union information. In the asymptotic regime, the leading term of the bias of the Union is equal to the sum of the leading term of the biases of single neuron information (see Eq. (S39) above). Thus the redundancy is expected to have relatively small bias. This prediction is also confirmed in all numerical simulations (Fig. 1, 2, S3, S4, S5, S6, S7, S8, S10, S11).

The unique information is the difference between the single neuron information and the redundancy. Given that the redundancy has little bias the unique information should be biased like the single cell information, which grows linearly with the number of discrete single cell responses. Thus the bias of the sum of the unique information should be similar to that of the Union and grow linearly with the number of single neuron responses. Thus, the unique information should have an intermediate bias between synergy and redundancy. Once again, these predictions are too confirmed in all numerical simulations (Fig. 1, 2, S3, S4, S5, S6, S7, S8, S10, S11).

In sum, the above explains why synergy is more biased than the other terms and gives an idea about the bias of the unique information and of the redundancy, suggesting that the bias of synergy is positive, the largest and grows quadratically with the number of single cell responses, the bias of the unique information is positive, the second largest and grows linearly with the number of single cell responses, and redundancy has the smallest bias which grows sub-linearly. It explains why the bias of synergy and joint get proportionally bigger for large numbers of possible discretized single cell responses.

Importantly, the analytical expression also can explain why the synergy bias of the shuffled data (and in general of the bias with lower information levels) are larger than the bias of the original distribution and give an idea of the situations in which they are expected to be tight. Take for example the joint information for which the bias is mostly dictated by the number of relevant response bins *R*_*J−s*_ of the stimulus-specific joint response distribution. Informative cases will have stimulus-specific distributions that are restricted to a fewer number of relevant bins than the total number of bins (because informative cases have less variability at fixed stimulus so more restricted and entropic stimulus-specific distributions) and also the bins that are relevant for a specific stimulus would not all be relevant for other stimulus-specific distributions (because informative cases will also show diversity of response distributions to different stimuli). When we randomly shuffle the trials combining trials to different stimuli, the shuffling operation will mix up responses and thus create larger stimulus-specific distributions (larger numbers of stimulus-specific relevant bins) which will then increase the bias, as found in our simulations. This prediction is confirmed by all our simulations. This result is important because it implies that bias corrections based on subtracting shuffled estimators lead a residual downward bias, which is useful to produce lower bounded or conservative estimates. It also implies that the shuffled-subtracted bias corrected estimates will be precise and not too conservative when the information level in the original data is low (because in this case the number of relevant bins of the shuffled and original data will be similar) and it will be too conservative when the information levels in the data are high (because in this case the number of relevant bins of the shuffled distribution will be much larger than the corresponding one for the original distribution). This leads to the design of tighter downward biased estimators of synergy that may be helpful to draw conservative yet sufficiently accurate conclusions.

While we derived these bias properties for the BROJA PID, we expect that the same conclusions would hold across PIDs. In particular, we would expect them to hold for the two other PIDs that we implemented, that is the *I*_*min*_ and the *I*_*MMI*_. We verified with simulations that these predictions indeed hold with *I*_*min*_ and the *I*_*MMI*_ (Fig. S6, S7, S10, S11). In fact in both these decompositions the Union information depends on the two marginal probabilities *p*(*r*_1_, *s*) and *p*(*r*_2_, *s*) (because they satisfy the so called pairwise marginals property). Thus all the considerations we made for the bias size of the BROJA Union are expected to hold also for the Union of these other definitions, because Union information definitions that satisfy the pairwise marginals property should have bias determined by the size of the finite size fluctuations of the marginals. The properties discussed for the BROJA held also for these other PIDs. If anything, we found that the bias of the synergy was even closer to the one of the joint information as the *I*_*min*_ and the *I*_*MMI*_ have the well documented problem of producing small values of unique information [19, 21].

### SM1.9 Venkatesh et al NeurIPS 2023 procedure for PID bias correction reformulated the discrete case

We tested the effectiveness in correcting for the bias of the PID bias correction procedures developed by Venkatesh and colleagues [18]. The procedure was formulated for Gaussian PID. We report it here for completeness, describing also the straightforward adaption of it that we did to use it in the discrete PID. In the following we use the subscript *N* to indicate the information values obtained by plugin of the empirical probabilities estimated from *N* trials s without using any bias correction, and (following Ref. [18]) we use the subscript *bias* − *corr* to indicate the information quantities corrected for the bias with the considered specific procedure.

The Venkatesh bias correction procedure [18] focuses first on the bias of the Union information. We consider union information *Union*(*S* : *R*_1_; *R*_2_) which is related to the other PID quantities from the following equation

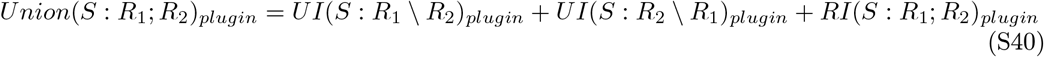

and is computed in the BROJA definition from Eq (6). In the following, we will indicate union information by omitting their dependency on source and target variables, as *Union*, for brevity. The Venkatesh procedure corrects for the bias by first computing the bias-corrected union information (see Eq. (17) and the rectification formulae Eq. (106, 107) of [18]) by computing the bias of the joint information with any of the method available for Shannon information (e.g. QE), then making the assumptions that it has the same bias as the joint information and rescaling it accordingly (Eq. (17) in [18]):

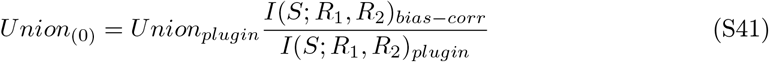

Note that, to follow precisely what was done by Venkatesh and colleagues [18], when applying the Venkatesh procedure if a Shannon information quantity had a negative value after the bias correction procedure, we set it to zero. Eq. (S41) assumes that the fraction of bias of the union information is the same as the one of joint information. As we found in our analytical calculations and simulations (Fig. S3), this assumption is incorrect, as union information is much less biased than joint information. Thus this first step will in most cases and for low trial numbers lead to a severe underestimation of union information. Then, to make sure all PID quantities after bias correction are non-negative and satisfy the PID linear constraints, the Venkatesh procedure applies a double post-hoc rectification of the bias-corrected value of the union information:

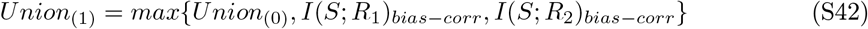

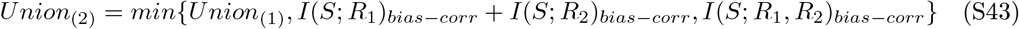

The resulting *Union*_(2)_ is the value we take in the following for bias corrected union *Union*_*bias−corr*_. From this the Venkatesh procedure computes the 4 bias-corrected PID terms using the bias-corrected union information and the bias corrected joint and individual information using the 4 linear PID constraints:

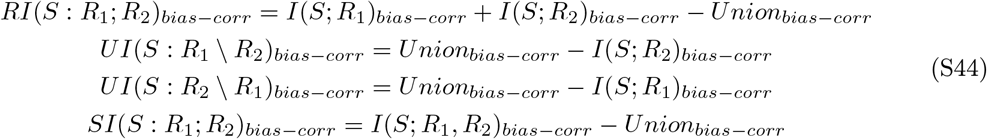

The problems of the Venkatesh procedure can be readily understood by considering scenarios in which both ground truth unique information components are larger than zero. Then, the major underestimation of *Union*_(0)_ in Eq. (S41) due to the incorrect assumption that the upward bias of the union information scales like the bias of the joint information (whereas in reality, as demonstrated in the previous section and in Fig. 1, S3, S4, S5, S6, S7, the former is small and scales linearly with the number of discretized single-neuron responses and the latter is large and scales quadratically) will lead to very low values of *Union*_(0)_. Then the first rectification in Eq. (S42) would take *Union*_(1)_ as the maximum between the information carried individually by the two sources, when in reality the unbiased union information should be larger than both single-source information, since both unique info are larger than zero. This would ultimately lead to a grossly underestimated *Union*_*bias−corr*_, which would result, from Eq. (S44) into an underestimation of the bias corrected unique information components (for which *Union*_*bias−corr*_ has a positive sign in Eq. (S44)) and an overestimation of the redundant and synergistic components (for which *Union*_*bias−corr*_ has a negative sign in Eq. (S44)). The major overestimation of synergy and redundancy and the major underestimation of unique information of the Venkatesh procedure have been consistently confirmed by our simulations (Fig. 2, 3, S8, S9, S10, S11).

### SM1.10 Details of experimental procedures for real neural data recorded from the mouse brain

#### SM1.10.1 Details of experimental procedures of mouse auditory cortex data recorded during a sound discrimination task

We reanalysed a previously published dataset [23] in which the activity of several tens to a few hundreds of neurons was recorded simultaneously using in vivo two photon calcium imaging from A1 L2/3 neurons in head-fixed transgenic mice during a pure-tone discrimination task (Fig. 4A). The experimental task was structured as follows. After a pre-stimulus interval of 1 s, head-fixed mice were exposed to either a low-frequency (7 or 9.9 kHz) or a high-frequency (14 or 19.8 kHz) tone for a period of 1 s. Mice were trained to report their perception of the sound stimulus by their behavioural choice, which consisted of licking a waterspout in the post-stimulus interval (0.5–3 s from stimulus onset) after hearing a low-frequency tone (target tones) and holding still after hearing high-frequency tones (non-target tones). Two-photon calcium imaging was used to continuously acquire the calcium fluorescence signals from individual A1 L2/3 neurons during the task with an imaging frame rate of 30 Hz. Calcium fluorescence traces were deconvolved as reported in [23] to estimate spike rates in each imaging frame. Full details of the experimental procedures and details of the ethical approval are reported in full in the original publication [23]. We analysed neurons recorded from *n* = 12 Fields of View (FOV) from *n* = 12 animals. For consistency with the previously published work [23] for the PID analysis reported in this paper (Fig. 4) we only considered individual neurons that carried significant intersection information [71], that is information about the stimuli that is used to inform the behavioral discrimination, according to the statistical permutation tests described in [23]. This led to selecting for PID analysis *n* = 375 individual neurons (out of the 2792 recorded neurons) leading to *n* = 6209 pairs of simultaneously recorded neurons used for the PID analysis.

For the PID analysis of these data, we identified for each neuron the imaging time frame within the trial of maximal intersection information [57] exactly as described in [23]. We then considered for each neuron a time frame of *n* = 10 imaging frames (corresponding to a window of 333 ms) around the peak intersection information time frame (we call this the peak time window for the neuron). Then we discretized activity for each neuron into *R* = 3 bins according to whether it was detected 0, 1 or *>* 1 spikes in the peak time window. The stimulus set used for this analysis was binary (*S* = 2), dividing the presented sound tones into the low- (*s* = 0) and high-frequency (*s* = 1) categories. For consistency with the original experimental publication [23], which analyzed joint stimulus information carried by pairs of neurons without breaking it up with the PID, we estimated the total stimulus information that was jointly carried by pairs of neurons following a time-lagged approach. Namely, we computed the time-lagged stimulus information carried jointly by the activity of each pair of neurons using Eq. (2) and using as responses *r*_1_ or *r*_2_ the discretized responses of each neuron measured as detailed above in their respective peak time windows.

#### SM1.10.2 Details of experimental procedures of mouse posterior parietal cortex data recorded during a decision-making task requiring sound localization

We analyzed previously published [53] neural recordings from a sound localization task in which mice reported perceptual decisions about the location of an auditory stimulus by navigating through a visual virtual reality T-maze (Fig. 4C). As mice ran down the T-stem, a sound cue was played from one of eight possible locations in head-centered coordinates. Mice reported whether the sound originated from their left or right by turning in that direction at the T-intersection. During each session, the activity of approximately 50 (range 37-69) layer 2/3 neurons was imaged simultaneously using two photon microscopy and the GCamp6f indicator. Calcium fluorescence traces were deconvolved as described in Ref. [53] to estimate spike rates in each imaging frame. We computed the mutual information between the joint activity of two simultaneously recorded neurons *r*_1_ and *r*_2_ and the stimulus category of the sound location (*S* = 2; in other words we computed information about whether the sound came from left or right of the midline, which is what the mice were asked to categorize in their task). This information is relevant for posterior parietal cortex, because this is an area that has been described as a sensory-motor interface that converts sensory information into signals relevant for decision-making. For the PID analysis of these data, for consistency with the above reported analysis of auditory cortex data, we identified for each neuron the imaging time frame within the trial of maximal stimulus information. We then considered for each neuron a time frame of *n* = 5 imaging frames (corresponding to a window of 320 ms) around the peak information time frame (we call this the peak time window for the neuron). Then we discretized activity for each neuron into *R* = 3 bins according to whether it was detected 0, 1 or *>* 1 spikes in the peak time window. Again, for consistency with the auditory cortex analysis, we estimated the total stimulus information that was jointly carried by pairs of neurons following a time-lagged approach. Namely, we computed the time-lagged stimulus information carried jointly by the activity of each pair of neurons using Eq. (2) and using as responses *r*_1_ or *r*_2_ the discretized responses of each neuron measured as detailed above in their respective peak time windows.

We analyzed neurons recorded from *n* = 11 Fields of View (FOV) from *n* = 11 animals. For consistency with the original publication reporting these data, we did not perform any selection of these neurons and we used all *n* = 713 neurons available in the published database, which allowed us to use *n* = 10750 pairs of simultaneously recorded neurons for the PID analysis.

#### SM1.10.3 Details of experimental procedures of mouse hippocampus data recorded during spatial navigation in a virtual environment

We reanalysed a previously published dataset [55] in which the activity of several tens to a few hundreds of neurons was recorded simultaneously using in-vivo two-photon calcium imaging from CA1 neurons in head-fixed transgenic mice during virtual reality navigation of a linear track (Fig. 4A). Calcium fluorescence signals were obtained using the jRCaMP1a calcium indicator and an imaging frame rate of 0.333*s*. Full details of the experimental procedures and details of the ethical approval are reported in full in the original publication [55].

We analyzed neurons recorded from *n* = 11 Fields of View (FOV) from *n* = 7 animals. Because hippocampal neurons encode position in space, we computed the mutual information between the spatial position *S* in which the animal was located in the linear track and the joint activity of two simultaneously recorded neurons *r*_1_ and *r*_2_. For consistency with the previous study reporting the original data [55], published study, the spatial position *s* along the linear track was computed by binning the space along the track into *S* = 12 equipopulated bins. For consistency with the previous study [55], to quantify the neural activity *r*_*i*_ of each neuron, we took the calcium traces and binned them into *R* = 2 equipopulated bins (only raw calcium traces and not deconvolved signals were available from Ref. [55]). For the PID analysis, we used all the *n* = 870 individual neurons present in the dataset leading to *n* = 36158 pairs of simultaneously recorded neurons used for the PID analysis.

### SM1.11 Simulated example of application of PID to within-network synergistic information transfer

**Figure S13:**
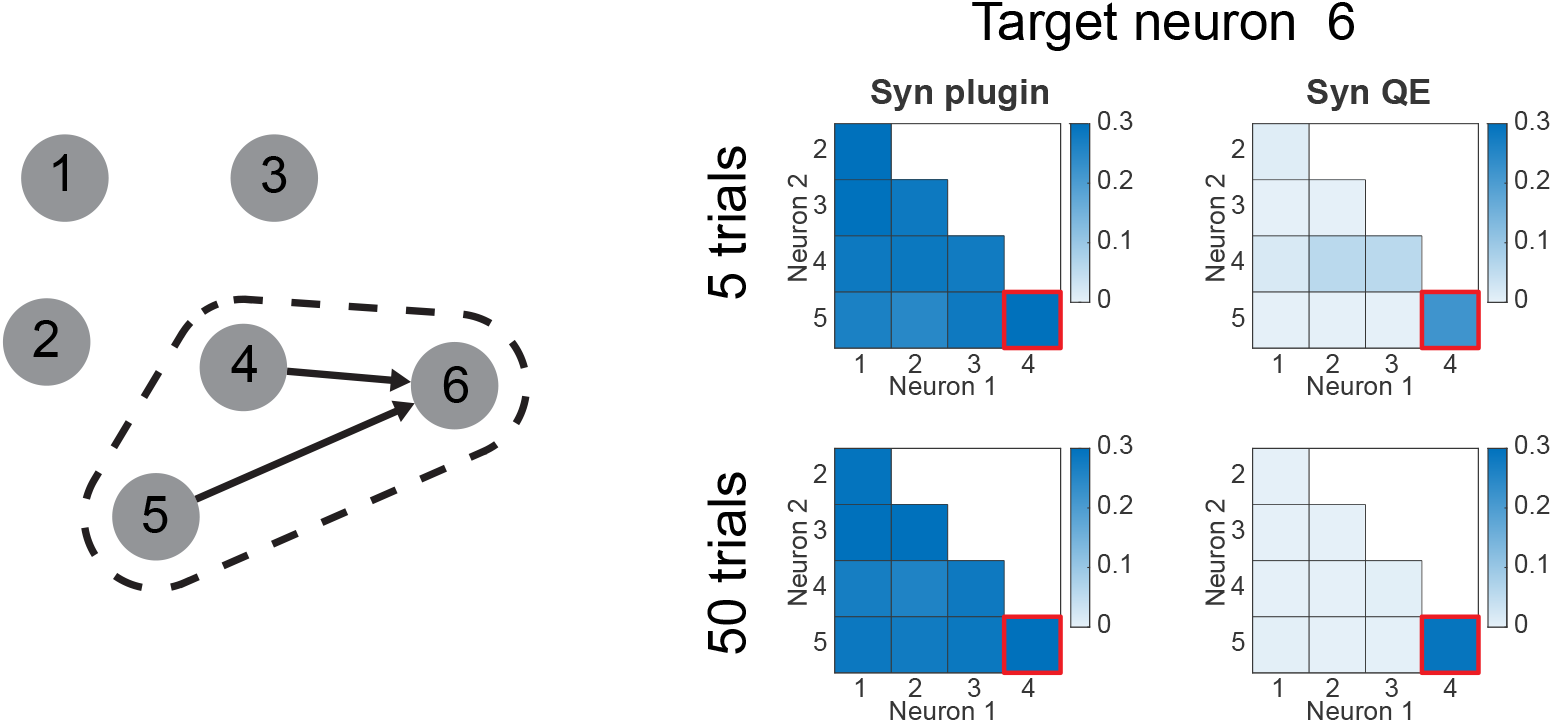
Example of using PID to describe triplet interactions in a network. On the left is shown a schematic of the network, where only neuron 6 is influenced by other neurons (neurons 4 and 5). On the right, the values obtained by setting neuron 6 as the PID target and going through all other possible cell pairs as sources. On the top row, the values are shown for the mean of 5 trials and on the bottom row, the mean across 50 trials. The left panels show the synergy values obtained naively without any bias correction. On the right panels, the corrected values using QE are shown. A red square points to the cell pair that has a true influence on cell 6.

To illustrate and test the usefulness of using PID to map information transfer within a network [22, 13, 13], we simulated a small network of neurons and computed the PID using the activity of another neuron (and not a sensory stimulus (*S*) as a target variable). We compared results using or not using sampling bias corrections.

We simulated a network (Fig. S13 of 6 nodes (to be conceptualized as different neurons or brain regions). Each node was modeled as Gaussian processes across time (100 timesteps). Nodes 1 to 5 are independent of each other with mean zero and unit standard deviation, while the activity of neuron 6 at time *t* is defined as the sum of the activities of neurons 4 and 5 at time *t* − 1 plus noise. Thus, the ground truth in this simulated dataset is that nodes 4 and 5 exert a synergistic information transmission on neuron 1, and there is no transmission of information between any other set of nodes. Thus, if we select neuron 6 as a target for our PID analysis, only the cell pair 4-5 should have significant synergy values [21].

The PID analysis was done using the BROJA redundancy measure [20], binning the neural activities with 4 equipopulated bins. The target was the activity of neuron 6 from time 2 to 100 and the sources were taken from the other cells from time 1 to 99 to account for the time-lagged nature of the interactions. The first row shows the result of running 5 trials of the same network and averaging its values. The second row shows the mean results of 50 trials.

The top left heatmap on Fig. S13 shows the synergy values for each possible cell pair targeting neuron 6 averaged across 5 trials. All the cell pairs have comparable, positive synergy values, which are significant according to a one-tailed t-test. When applying a QE correction of one instantiation of the simulation, only the pair corresponding to the real connections is still significantly above zero (one-tailed t-test *p <* 0.01). When increasing the number of simulations to 50, the plugin PID is still unable to assign a significant synergy value only to the real source neurons (Fig. S13 bottom left, all pairs have significant values with *p <* 0.001). After the QE correction, only the real cell pair remains significantly above zero (one-tailed t-test *p <* 0.001).

Together, these results show that the bias corrections are useful also to obtain better estimates of information about target variables being the activity of other neurons (as often done in neuroscience [22, 13, 13]) and that the bias corrections are useful to identify patterns of synergy within larger networks.

### SM1.12 Data and code availability

Code for simulating and analysis is available from the corresponding author upon reasonable request. The real neural data used here in Fig. 4 were published before. Data of [23] were released with the original publication and can be found at https://doi.org/10.1101/2021.08.31.458395. Data of [53] were republished in [72] and can be found at https://doi.org/10.12751/g-node.tqbad8. Data of [55] are available from the corresponding author upon reasonable request.

